# Using high abundance proteins as guides for fast and effective peptide/protein identification from metaproteomic data

**DOI:** 10.1101/2020.05.21.109348

**Authors:** Moses H. Stamboulian, Sujun Li, Yuzhen Ye

**Affiliations:** Luddy School of Informatics, Computing and Engineering, Indiana University, 700 N. Woodlawn Avenue, Bloomington, IN 47408

**Keywords:** high abundance protein (HAP), sample profiling, expanded target database, spectra search, human gut metaproteomics

## Abstract

**Background:** A few recent large efforts significantly expanded the collection of human-associated bacterial genomes, which now contains thousands of entities including reference complete/draft genomes and metagenome assembled genomes (MAGs). These genomes provide useful resource for studying the functionality of the human-associated microbiome and their relationship with human health and diseases. One application of these genomes is to provide a universal reference for database search in metaproteomic studies, when matched metagenomic/metatranscriptomic data are unavailable. However, a greater collection of reference genomes may not necessarily result in better peptide/protein identification because the increase of search space often leads to fewer spectrum-peptide matches, not to mention the drastic increase of computation time.

**Methods:** Here, we present a new approach that uses two steps to optimize the use of the reference genomes and MAGs as the universal reference for human gut metaproteomic MS/MS data analysis. The first step is to use only the High Abundance Proteins (HAPs) (i.e., ribosomal proteins and elongation factors) for metaproteomic MS/MS database search and, based on the identification results, to derive the taxonomic composition of the underlying microbial community. The second step is to expand the search database by including all proteins from identified abundant species. We call our approach HAPiID (HAPs guided metaproteomics IDentification).

**Results:** We tested our approach using human gut metaproteomic datasets from a previous study and compared it to the state-of-the-art reference database search method MetaPro-IQ for metaproteomic identification in studying human gut microbiota. Our results show that our two-steps method not only performed significantly faster but also was able to identify more peptides. We further demonstrated the application of HAPiID to revealing protein profiles of individual human-associated bacterial species, one or a few species at a time, using metaproteomic data.

**Conclusions:** The HAP guided profiling approach presents a novel effective way for constructing target database for metaproteomic data analysis. The HAPiID pipeline built upon this approach provides a universal tool for analyzing human gut-associated metaproteomic data.

## Introduction

Culture independent studies of microbial communities associated with different environments are promoted by two main reasons: the discovery of the growing importance of these communities contributing to their environment/host, and the rapid advancements in sequencing technologies [1, 2, 3, 4]. Of these communities a particular attention has been devoted to the human gut microbiota for its impacts on human health and diseases [5, 6, 7] and its potential applications to improving the efficacy of treatments (including cancer chemotherapy and immunotherapy) [8, 9], and prevention of diseases (e.g., using probiotics [10, 7]). Numerous studies, focusing on human gut microbiota, have already been conducted showing its central role in regulating human health, reporting the latter’s deterioration to be directly related to dysbiosis in the composition and functionality of gut bacteria [11]. Irritable Bowel Syndrome (IBS), Inflammatory Bowel Diseases (IBD) and *Clostridium difficile* Infection (CDI), just to mention few, are examples of diseases that are found to be associated with the imbalance within the human gut microbiota [12, 13, 14]. It has also been shown that the genetic makeup and the diet of the host have direct impacts on the composition of the gut bacteria, while in the meantime the latter regulating digestive and metabolic (and beyond) processes of the host, creating a symbiotic relationships between the two [15, 16, 17].

Improvements of both the experimental techniques (e.g., sequencing technology and sample collection [18]) and computational methods (such as those for binning and assembly [19]) have accelerated the microbiome research. While metagenomics and metatranscriptomics are essential for quantification and characterization of taxonomic compositions of microbial communities, they only suggest possible metabolic potential and are unable to confirm the actual presence of such biological processes in the communities, since most biological functions are carried out at the protein level. Shotgun proteomics, which studies all the translated proteins in a sample recovered from the environment directly, has been shown to be promising in uncovering functional information about gut bacteria [20, 21, 22]. Combining information from multiple-omic experiments will provide opportunities for more comprehensive characterization of the functionalities of the underlying microbial communities.

The initial shotgun metaproteomic experiments date back as far as a decade ago [20], and despite the numerous improvements of the employed technologies ever since, metaproteomics still relies on mass spectrometers that are mainly designed to handle single species samples at a time. Unlike sequencing technologies, shotgun metaproteomics still suffers from the diversity and complexity of microbiome communities, making it challenging for data evaluation and downstream analysis [23, 24]. A typical metaproteomic data analysis includes these steps: construction of a sample-specific target protein sequence database, peptide identification against the target database, and downstream functional analysis [25]. Without any previous knowledge concerning the active organisms prominent in the target sample, the results and the quality of downstream analysis highly depend on the constructed protein database [26]. Using large and expanded database to include a comprehensive set of species for spectral search may reduce the search sensitivity, making it difficult to estimate false discovery rate (FDR) without the expense of increased false negatives while significantly increasing the search time [27]. On the other hand, manually constructing a customized target database isn’t straightforward, given the complexity and diversity of the human gut flora, and thus is not commonly adopted in practice. In the cases when multi-omics datasets are available for the same microbial community, the metagenomic and/or metatranscriptomic data can be utilized to derive the target protein database for metaproteomic data analysis. To reduce cost and time often one representative sample is sequenced and the derived target database is shared among the subsequent metaproteomic studies [28, 29]. We have previously developed novel algorithms (Graph2Pro [30] and Var2Pep [31]) to optimize the use of matched metagenomic/metatranscriptomic data for metaproteomic data analysis.

Many metaproteomic datasets have been and will be produced without matched metagenomic sequences information, so it is important to develop methods for analyzing these metaproteomic datasets without matched metagenomic databases. In addition, it is attractive to develop a universal framework for metaproteomic data analysis across different samples and studies without relying on specific metagenomic datasets. The success of such universal approaches relies on 1) the availability of a comprehensive reference protein database for spectral search; and 2) algorithms that enable effective use of the large reference database. Microbiome research has drastically expanded the protein universe related to microbial species. On the other hand, the two-step method, which uses matches derived from a primary search against a large database to create a smaller subset database for false discover rate (FDR) controlled second step search, has shown improved sensitivity in peptide identification from metaproteomic data, as shown in Jagtap et al. [27]. Zhang et al. [32] developed MetaPro-IQ, which leverages the large gene catalogues (for human gut microbiome and mouse gut microbiome) as the target databases in its first step of spectra searches. MetaPro-IQ identified 15,200 peptides on average over samples collected from intestines of eight human subjects, matching the results using matched metagenome approach on the same datasets. MetaPro-IQ was later integrated into an automated pipeline MataLab [33], which also leverages spectra clustering to improve the speed of peptide identification from database searches.

Spectral search against a large reference database (as in MetaPro-IQ) is computationally intensive. Taking advantage of the recent expansion of the human gut microbial genomes [34, 35], we developed a new two-step approach for metaproteomics data analysis, using over 3,000 reference genomes and MAGs by first profiling microbial communities based on the spectral search against a database of high abundance proteins (HAPs) encoded by these genomes. The profiling results are then used to guide the construction of the target database for the second step spectral search, including all putative proteins encoded by only the prominent genomes identified in the first step. As a result, our approach significantly reduces the computational cost of the whole process. We call our approach HAP guided Metaproteomics IDentification or simply HAPiID (pronounced as: Happy ID). We tested HAPiID using eight publicly available metaproteomic datasets [32]. The results show that HAPiID outperformed MetaPro-IQ in both the number of identified peptides and the speed. We note that in this paper we compared HAPiID with MetaPro-IQ [32] (instead of MetaLab [33], which uses MetaPro-IQ for peptide identification), to emphasize the promise of developing novel approaches for constructing effective target database for spectra searches in metaproteomic studies.

## Materials and Methods

### The overall approach and the rationale

HAPiID uses two steps for metaproteomic MS/MS data identification. The first step is to infer the taxonomic profile of microbial community based on metaproteomic data, by searching spectra against a database containing only the proteins that are likely to be highly abundant due to their functional importance in any species. In this step, HAPs from all gut reference genomes and MAGs are considered. The second step is to do spectral search against an expanded database of all proteins but from a much smaller selection of reference genomes with most spectral support according to search results from the first step. The rationale of the two-steps approach and using highly expressed proteins in the first step is that if these high abundance proteins (which are highly conserved and are of important functions to any microbial species) encoded by a genome are not detected by metaproteomics approach, other proteins encoded by the same genome are less likely to be detected. The purpose of the first step is two folds: 1) to profile a metaproteomic sample and identify species prominent to it, and 2) to expand these prominent species to construct a sample-specific target protein database for subsequent peptide identification.

### Gut reference microbial genomes/MAGs

To assemble reference genomes for MS/MS identification in human gut metaproteomics, we collected genomes from two recent studies [34], [35]. Bacterial genomes reported in [35] were compiled from two sources: a total of 617 genomes obtained from the human microbiome project (HMP) [36], and 737 whole genome-sequenced bacterial isolates, representing the Human Gastrointestinal Bacteria Culture Collection (HBC). These 737 bacterial genomes were assembled by culturing and purifying bacterial isolates of 20 fecal samples originating from different individuals [35]. The bacterial genomes reported in [34] were generated and classified from a total of 92,143 metagenome assembled genomes (MAGs), among which a total of 1,952 binned genomes were characterized as non-overlapping with bacterial genomes reported. These novel binned genomes were termed as Uncharaterised MetaGenome Species (UMGS). We also amended 56 archaeal genomes belonging to 13 species that were shown to be essential inhabitants of the human digestive track [37].

We were able to retrieve 612 out of 617 RefSeq sequences using the reported RefSeq IDs. Our final dataset for this study contains 612 genomes from the RefSeq database, 737 whole genome-sequenced bacterial isolates from the HBC dataset, 1,952 UMGS genomes and 56 archaeal genomes, making a total of 3,357 genomes and MAGs.

We applied the least common ancestors approach GTDBTK [38] to assign taxonomic labels to these genomes. This approach was able to assign *order* level taxonomies to 3,293 out of these genomes.

### Identification of ribosomal proteins and elongation factors (HAPs)

We first collect (from RefSeq genomes) or predict (from MAGs) putative proteins and then identify highly abundant proteins, i.e., ribosomal proteins and elongation factors among them. FragGeneScan [39] was employed, with default settings to predict protein coding genes and their respective amino acid sequences, from MAGs. A total of 2,602,889 genes were predicted from the HBC contigs, and another 4,001,749 genes from the UMGS bins. Genes for the RefSeq genomes were obtained from the RefSeq database. A total of 2,017,525 genes were downloaded for all the RefSeq genomes used. The final dataset contains over eight million putative proteins.

We extracted ribosomal proteins and elongation factors from the 612 RefSeq annotated genes, by searching for keywords “ribosomal protein” and “elongation factor”, then we scanned these genes against the Pfam database (Pfam32.0) [40], using hmmer3 program [41] to extract the Pfam profiles that confidently match with these sequences. A strict E-value cutoff of *e*^−10^ was used to report hits. A total of 120 Pfam profiles had significant hits with the Ref-Seq sequences; however some of these domains were irrelevant (i.e. tRNA synthetase), and were present due to their co-presence with relevant domains in multi-domain proteins. After manually curating these profiles, we kept domains that were only for ribosomal proteins and elongation factors. A final list of 77 Pfam profiles were retained (the list is included in the HAPiID pipeline). All putative proteins predicted from the HBC and UMGS bins were then scanned against these Pfam profiles using hmmscan [41], to identify ribosomal proteins and elongation factors. A total of 39,584, 48,598, 101,899 and 2,074 ribosomal proteins and elongation factors were extracted from RefSeq genomes, HBC bins, UMGS bins and archaeal genomes, respectively. We note that all of the 3,357 gut genomes have these highly expressed proteins, ranging from 7 to 80 HAPs. A more detailed distribution of the number of marker genes across the different genome sources is summarized in Supplementary Figure S1. After removing redundant proteins (with 100% sequence identity according to CD-HIT [42]), we constructed the *High Abundance Protein database* (*HAPdb*), which contains 110,103 proteins in total, to be used as the target database for the first step search in HAPiID pipeline.

### HAPiID pipeline

Figure 1 shows the overall structure of the HAPiID pipeline. The first step is the *sample profiling*, which involves searching MS/MS data against HAPdb to profile the sample of interest. Peptides identified in this step are used to quantify the presence of prominent species. We implemented a simple greedy approach that reports the minimum list of genomes needed to cover all identified spectra. The greedy approach works by first ranking all the genomes in decreasing order of the number of identified unique spectra they can explain, and then choosing the top *n* most abundant genomes to construct expanded target database (for the second-step search). In principle, *n* is chosen such that all spectra are covered, which however doesn’t work in practice due to the existence of a large number of genomes with only a few spectra (which could also be false identifications). Instead, we devised an automatic procedure for choosing the parameter *n* such that these *n* genomes cover at least 80% (or a user defined %) of the identified spectra in the profiling phase, which worked well in practice (see Results).

**Figure 1:**
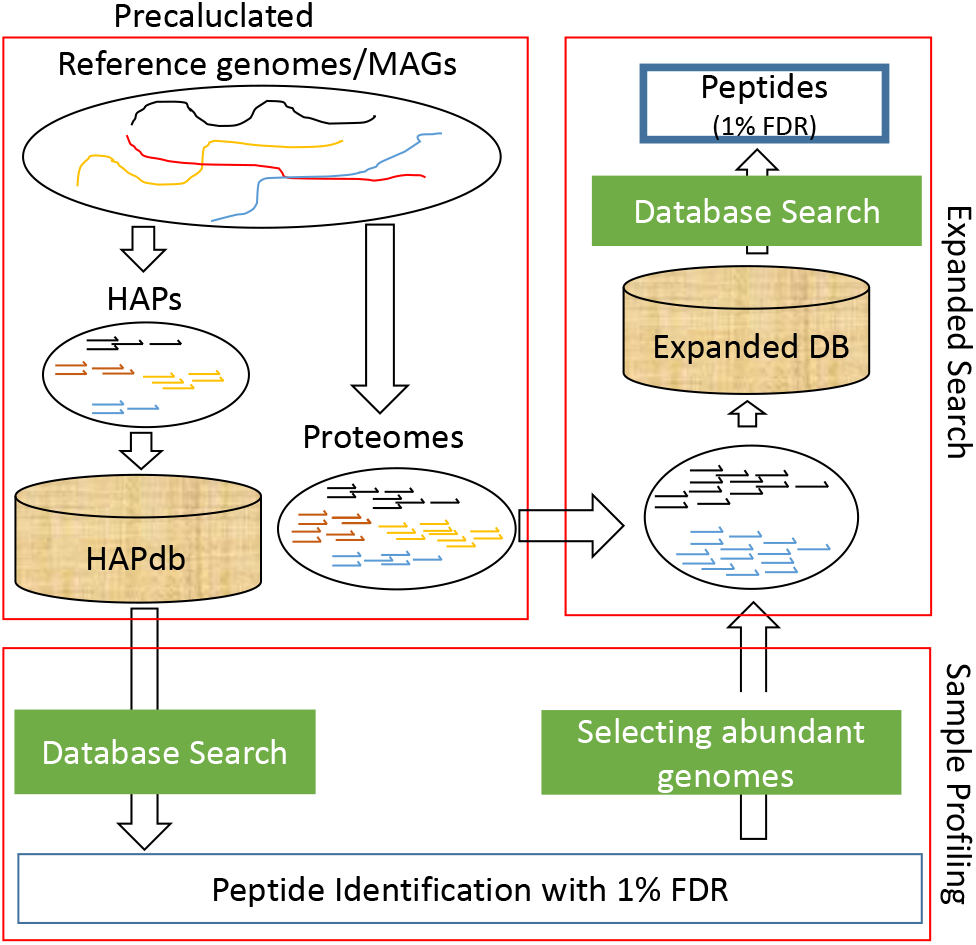
An overview of the HAPiID pipeline for peptide identification from metaproteomic MS/MS data. The pipeline is composed of two main steps, *sample profiling* (which is based on searching spectra against precalculated HAPdb containing HAPs from all genomes/MAGs) and *expanded search* (to search spectra against all proteins from a smaller selection of abundant genomes).

The second step is the *expanded search*, where the spectra are searched against the expanded target database constructed based on the sample profiling results. Species diversity and composition will also be reported at the end of the search.

We tested both MS-GF+ [43] and X! Tandem [44] as the search engines for identifying tandem mass spectra using a target protein database. However, our pipeline can be modified to work with other search engines. In both steps (profiling and targeted search), a strict FDR of 1%, which is commonly adopted, was used to filter out potential false positives from our final peptide identification. The FDR was estimated using the target-decoy approach [45], where the reverse protein sequences were used as the decoys.

### Metaproteomic datasets

We tested our pipeline using one synthetic and eight publicly available human gut metaproteomic datasets [46, 32]. The synthetic dataset called SImplified HUman Interstinal MIcrobiota (SIHUMI) was produced from proteomes of eight genomes (*Anaerostipes caccae, Bacteroides thetaiotaomicron, Bifidobacterium longum, Blautia producta, Clostridium butyricum, Clostridium ramosum, Escherichia coli*, and *Lactobacillus plantarum*) [46]. The later eight metaproteomic datasets were obtained from children all under 18 years old during colonoscopy from children’s hospital of Eastern Ontario. The use of these datasets will facilitate the comparison of our approach to MetaPro-IQ and results reported from their matched metagenome approach as well [32].

We tried both search engines when applying HAPiID to these datasets. We used the MS-GF+ search engine (version v10089) [43] with the following parameters: high-resolution LTQ (instrument type), precursor mass tolerance of 15 ppm, −1-2 for the isotope error range, allowing at most 3 modifications including variable oxidation of methionine and fixed carboamidomethy of cysteine, maximum charge of 7, minimum charge of 1, and allowing semi-tryptic fragmentation. We used the same parameters for X! Tandem (VENGEANCE 2015.12.15) as reported in MetaPro-IQ [32]: up to two miss-cleavages (trypsin/P), carbamidomethylation of cysteine as a fixed modification, oxidation of methionine as a potential modification, a fragment ion tolerance of 20 ppm and a parent ion tolerance of 10 ppm.

### Functional annotation of identified proteins

We used two sources to assign functions to the proteins identified from metaproteomic data by HAPiID. The first one is KofamKOALA [47], which is based on KOfam, a customized database of KEGG orthologs [48]. The other one is Pfam database [49]. Both of these sources rely on HMMER tools to scan protein sequences against their databases for functional annotation [41].

### Availability of the pipeline

The HAPiID pipeline and the data required to use the pipeline are available as open source at https://github.com/mgtools/HAPiID.

## Results

We first evaluated the efficiency and accuracy of HAPiID using a synthetic metaproteomic dataset and eight real gut metaproteomic datasets. We then compared the performance of our pipeline to MetaPro-IQ. Finally we demonstrated the applications of our pineline including metaproteomics-based taxonomic profiling and studying the functional distribution of expressed proteins from highly abundant species using metaproteomic data.

### Evaluation of HAPiID using the synthetic gut metaproteomic dataset

Instead of searching spectra against more than eight million proteins predicted from the entire collection of 3,357 gut genomes, HAPiID involves two searches against much smaller databases: the first database contains only HAPs from all the genomes, and the second one contains all proteins predicted from a smaller collection of genomes. Selection of the genomes (based on the first profiling step) determines the efficiency and accuracy of the overall peptide identification by HAPiID. Here we use the synthetic SIHUMI dataset, for which we know the underlying genomes (see methods for more details), to evaluate the accuracy of the profiling step of the HAPiID and to learn about how to select genomes for the second step. We constructed an exact reference database consisting of proteins encoded by the eight genomes (called SIHUMI DB) for peptide spectra search using MS-GF+. The identified peptides were then used for estimating the species composition of the synthetic metaproteomic dataset. Figure 2(a) summarizes the species composition, which uses the percentage of unique spectra mapped to the underlying genomes to approximate the species abundance. Although the SIHUMI dataset was constructed using eight species, there were no unique peptides/spectra identified to *Lactobacillus plantarum*. In addition, the top five most abundant species (*Bacteroides thetaiotaomicron*, *Blautia producta, Anaerostipes caccae, Escherichia coli*, and *Clostridium ramosum*) accounted for 99.67% of identified spectra.

**Figure 2:**
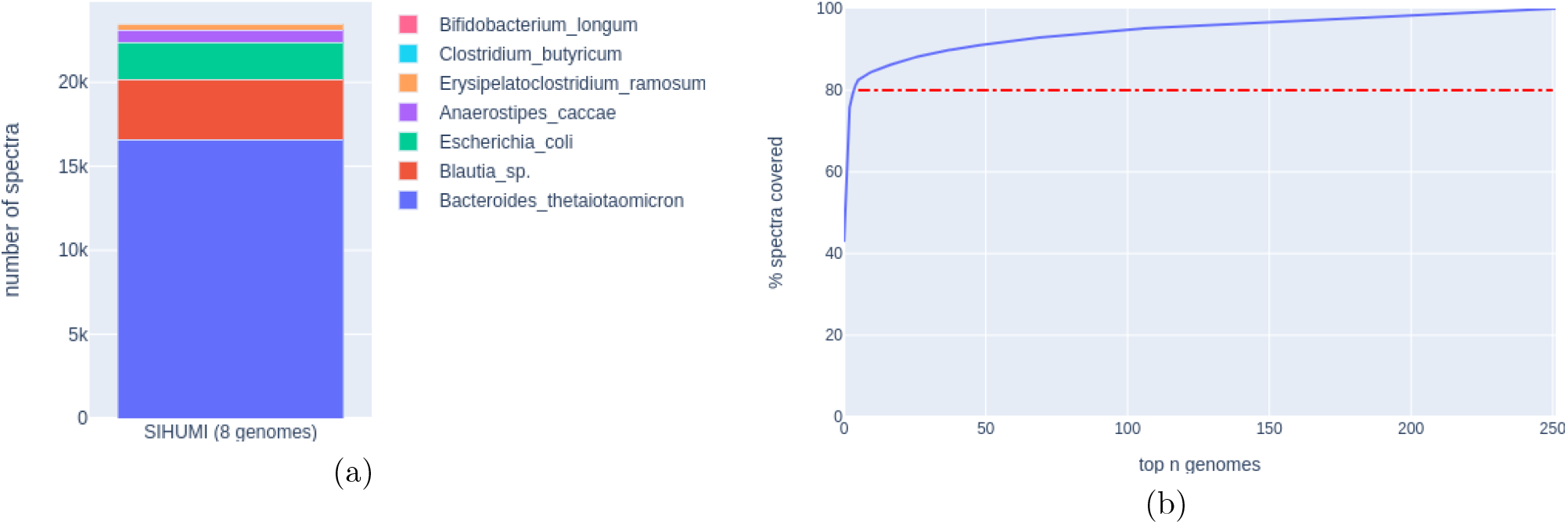
Test of HAPiID pipeline using the SIHUMI synthetic metagenomic dataset. (a) shows the relative abundance of the underlying species based on spectral identification using the eight underlying genomes as the reference database. (B) shows the percentage of unique spectra identified as a function of number of genomes selected by HAPiID.

Figure 2(b) shows the percentage of identified spectra as a function of top *n* species included during the profiling phase by HAPiID applied to the synthetic dataset. The plot shows that only the first few genomes contributed significantly to the identification of spectra: 80% of the identified spectra in the profiling step can be explained by the first five species, and after that only a very small fraction of spectra can be explained by including yet another genome. More importantly, we show that the top five genomes identified by HAPiD’s profiling step are the same as those revealed by the targeted search (against SIHUMI DB), and are in the same order as expected when ranked by their relative abundances. We note that the two species that were missed by HAPiID (*Bifidobacterium longum, Lactobacillus plantarum*)—when only genomes that cover at least 80% of identified spectra were included (this criterion worked well for the real gut metaproteomic datasets as well, as shown below)—contribute less than 0.4% of the total number of identified spectra. It would be difficult to identify peptides from such very rare species without increasing false identification, considering that HAPiID uses a large collection of genomes for the profiling step, necessary for its application to real metaproteomic datasets with unknown and much more complex species composition.

### Test of HAPiID using eight gut metaproteomic datasets

Next we benchmarked the efficiency of the HAPiID using more complex gut metaproteomic datasets. After the first profiling step, we ran HAPiID by selecting different numbers of genomes for the second step expanded search, and compared the results from the different runs. Figure 3(b) summarizes the total number of identified peptides when the second search used target databases containing putative proteins encoded by the top n (n = 5, 10, 20, 30, 40, 50, 100, 200) genomes. Results from the first step search (against HAPs only) are also shown, as a baseline to quantify the improvement in the identification rates when the pipeline expands to include all proteins encoded by each selected genome. The figure shows consistent increase of identified peptides when the number of whole genomes increases. The number of peptides remains roughly flat between 20 and 50 genomes, and the performance started to deteriorate when including more than 50 genomes, indicating that after this point, including more genomes will unnecessarily increase the search space that worsens the spectral identification.

**Figure 3:**
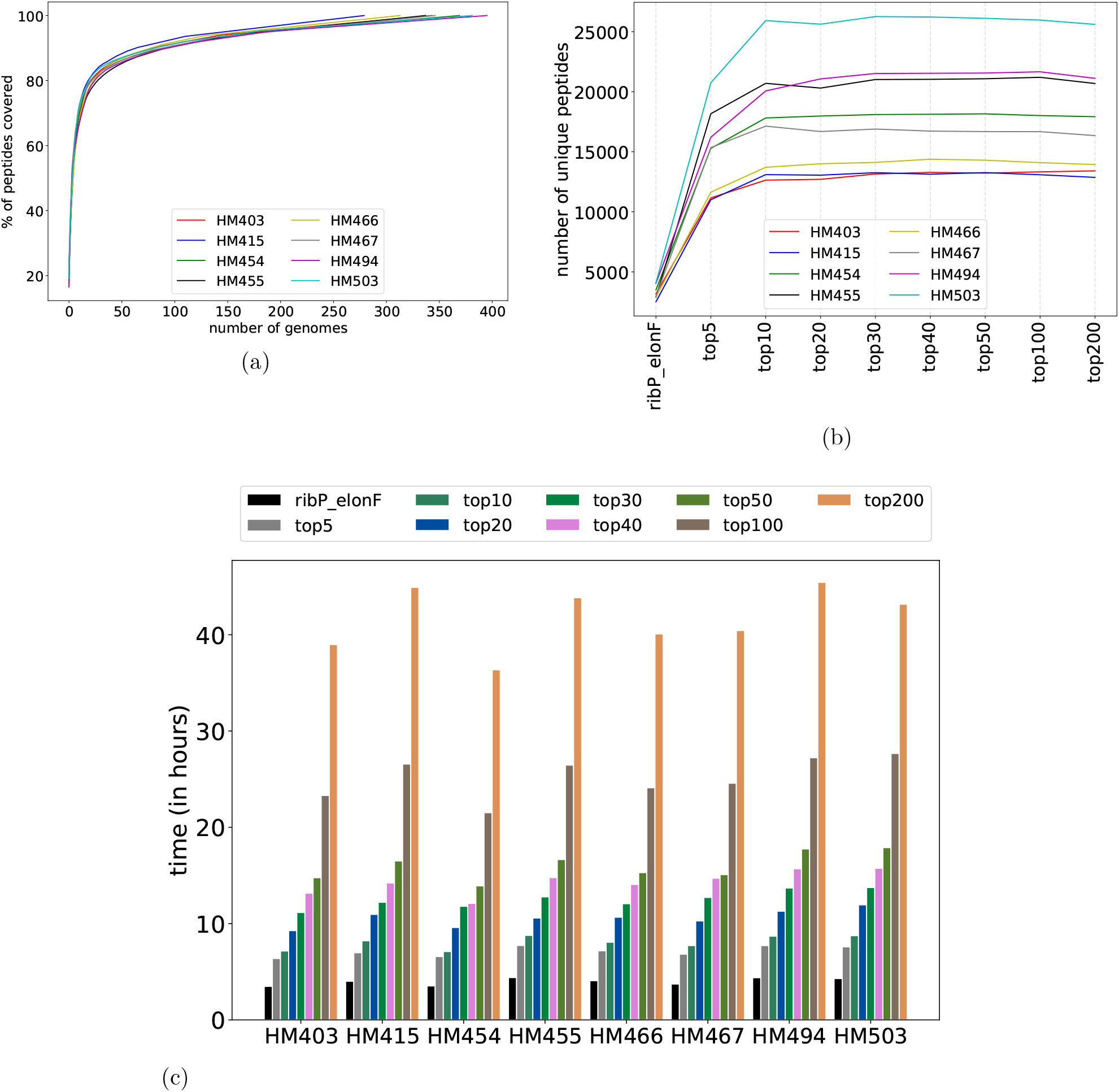
Test of HAPiID pipeline on eight human gut metaproteomic datasets. (a) and (b) show the number of unique peptides identified during profiling step, and number of unique peptides identified during expanded search, respectively. In (b), the ribP_elonF represents the results from the profiling step. (c) shows the running time (in hours) as a function of number of genomes included in the second step, expanded search.

A potential problem of using small target database for spectra search is that a spectrum may be identified as a wrong peptide because the true peptide, which can be identified when a larger target database is used, is not contained in the small target database. To address this problem (and to determine the appropriate size of the databases for the second step search for fast yet accurate peptide identification), we checked whether or not the same peptide is identified from the same spectrum when using a small or a big target database. For quantification purposes, we defined the *consistency rate* as the fraction of peptide-spectrum matches (PSMs) that remain the same when the size of the database was increased. We note that when comparing consistency rates across two databases, the small database is always a subset of the big database. Table 1 summarizes the average consistency of the peptide identification when databases of different sizes are used (in the second step search) for all eight metaproteomic datasets (results for individual samples are shown in Supplementary Tables S1 and S2). The results suggest that databases built from fewer than 20 genomes are not sufficiently large to produce accurate identifications; for example, peptide identifications based on top five genomes and top 100 genomes only had 97.8% agreement (i.e., the discrepancy is 2.2%, which is greater than the commonly used 1% FDR). However, when the number of genomes reaches 20 or more, the search results had about 99% agreement with the results based on searches against an expanded database built from for example, 100 genomes.

**Table 1:**
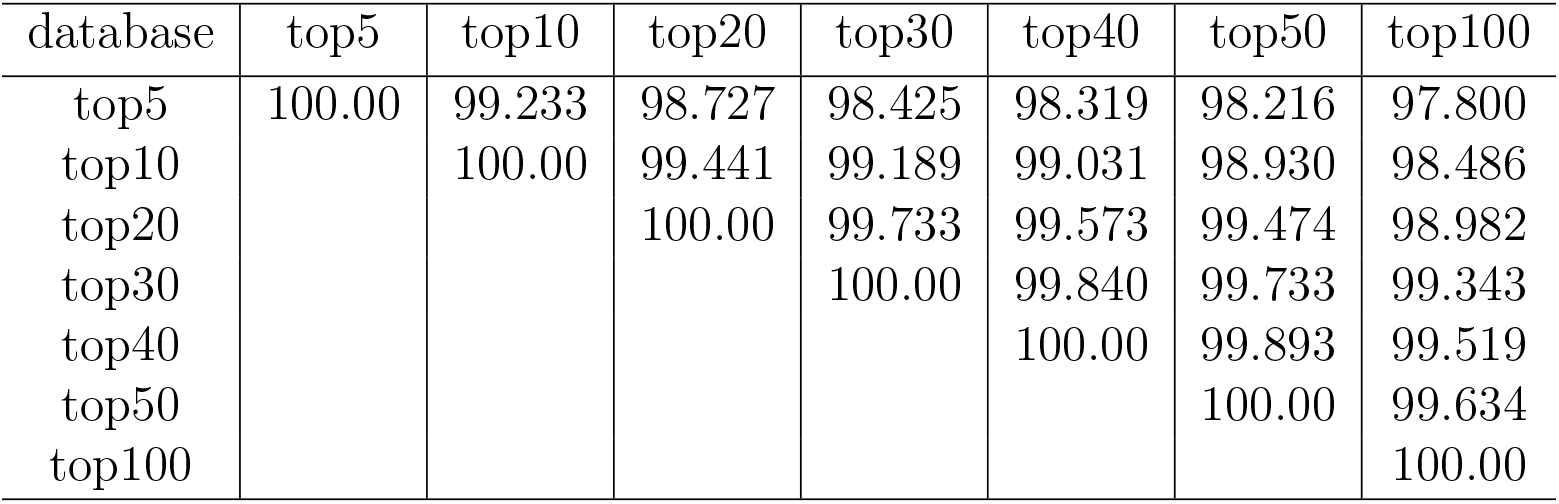
Agreement of identified peptides between searches against target databases of various sizes.

On the other hand, using fewer genomes speeds up the spectral search process. Figure 3(c) shows the running time (in CPU hours) of HAPiID using target databases of various sizes. Just as expected, the running time grows (linearly) with the number of genomes used for constructing the expanded search database. When using top 20 genomes in the second step of search, the whole pipeline (using MS-GF+ search engine) was finished in less than 11 hours for all the datasets we tested. This is a significant achievement, considering that speed is one of the major concerns about metaproteomic data analysis [43].

We also tested HAPiID using X! Tandem [50, 51], which was used in MetaPro-IQ [32] for its first and second steps of spectra match. We used the same parameters for X! Tandem as reported by MetaPro-IQ (see Methods). Table 2 summaries the peptide identification results by HAPiID with the two search engines. Overall, HAPiID using the two engines achieved comparable performances across different samples, with HAPiID using MS-GF+ marginally outperformed HAPiID using X! Tandem in six out of the eight cases. We also summarize the overlap of the identified peptides in Supplementary Figure S2.

**Table 2:** Comparison of the number of peptides identified by different approaches across the eight human gut metaproteomic datasets.

|  | HM403 | HM415 | HM454 | HM455 | HM466 | HM467 | HM494 | HM503 |
| --- | --- | --- | --- | --- | --- | --- | --- | --- |
| HAPiID-MSGF | 12,962 | **12,535** | **17,619** | 20,803 | **13,862** | 15,924 | **21,108** | **24,962** |
| HAPiID-X! | 12,414 | 12,180 | 17,314 | **21,145** | 13,676 | **16,334** | 21,042 | 24,459 |
| MetaPro-IQ | 12,606 | 11,562 | 15,446 | 17,868 | 11,302 | 12,380 | 18,562 | 21,879 |
| Matched | **13,156\*** | 12,179 | 15,863 | 18,677 | 11,733 | 12,724 | 19,248 | 23,632 |

\* The highest numbers of peptides identified in each sample are highlighted in bold; HAPiID-MSGF (HAPiID using MS-GF+ as the search engine); HAPiID-X! (HAPiID using X! Tandem as the search engine); Matched: peptide identification using matched metagenome as the reference. Results for the MetaPro-IQ and matched metagenome approach were taken from [32].

Considering all (the peptide identification efficiency as shown in Figure 3(b), the accuracy of the identification as summarized in Table 1, and the running time as shown in Figure 3(c)), using top *n* most abundant genomes that cover up to 80% of the total number of spectra during profiling phase, appears to be a good practice for the second step of expanded search for analyzing the eight human gut metaproteomic datasets. Using this criterion, on average 20 genomes were selected for expanded database search when tested over these eight datasets. We used the results based on this setting for the downstream analyses reported below.

### Comparison with MetaPro-IQ and matched metagenome approach

We first compared the peptide identification results from HAPiID (using MS-GF+ as the search engine) and MetaPro-IQ. Because the MetaPro-IQ pipeline was not publicly available, we used their reported identification results [32] for comparison. For the eight gut metaproteomic datasets we have tested, HAPiID method identified 17,472 peptides per sample on average, which is significantly higher than the results reported by MetaPro-IQ. Figure 4 summaries the peptide identification results for both methods and their overlap (see details of the comparison in supplementary Table S3). We note that we don’t distinguish Leu and Ile when comparing peptides as they are indistinguishable by mass spectrometry. In all eight samples we can see a significant overlap between the peptides identified by both approaches. However, HAPiID was able to identify significantly more peptides than MetaPro-IQ. On average there was 54% (11,519 peptides) overlap between peptides identified by both methods, while around 14% (3,683 peptides) of all the peptides were identified by MetaPro-IQ only, and more than 27% (5,824, peptides) of all the peptides were identified by our approach only, over all eight samples. We further examined the list of peptides only identified by MetaPro-IQ (3,683 peptides on average). Among them, around 40% (1218 peptides on average) were present in our target database, however they were identified with scores lower than the thresholds to pass the 1% FDR filtering.

It was shown in [32] that MetaPro-IQ achieved comparable performance as the spectral search using a matched metagenome to prepare search database for spectral match. By contrast, HAPiID resulted in identification of more spectra than the matched metagenome approach for seven out of the eight cases. Table 2 and Supplementary Figure S3 show the details of the comparison (Supplementary Figure S4 shows three way comparison). Combining all eight samples, a total of 29,074 unique peptides were identified by HAPiID but not the matched metagenome approach. We show that about 70% (20,342 peptides) of these HAPiID-only peptides could be explained by the top 50 genomes contributing to the identified peptides from the matched metagenome approach (since the matched metagenome approach didn’t provide species identification, we mapped its identified peptides onto HAPiID’s gut genome collection to reveal the possible underlying species). This result suggests that although matched metagenome approach provides more targeted reference database for metaproteomic data analysis, the reference protein database constructed from metagenome is likely incomplete (some proteins are missing due to the incompleteness of metagenome assemblies) and therefore making it less ideal for spectral match in metaproteomic data analysis.

We then compared the running time of MetaPro-IQ and HAPiID. Since MetaPro-IQ is not publicly available, we could not benchmark its execution time directly. Considering that the first step in MetaPro-IQ (searching spectra against the whole gut microbial gene catalog) is the computationally most demanding step (the second step search involves a reduced database), we focused on comparing the running time of our approach with the first step of MetaPro-IQ.

**Figure 4:**
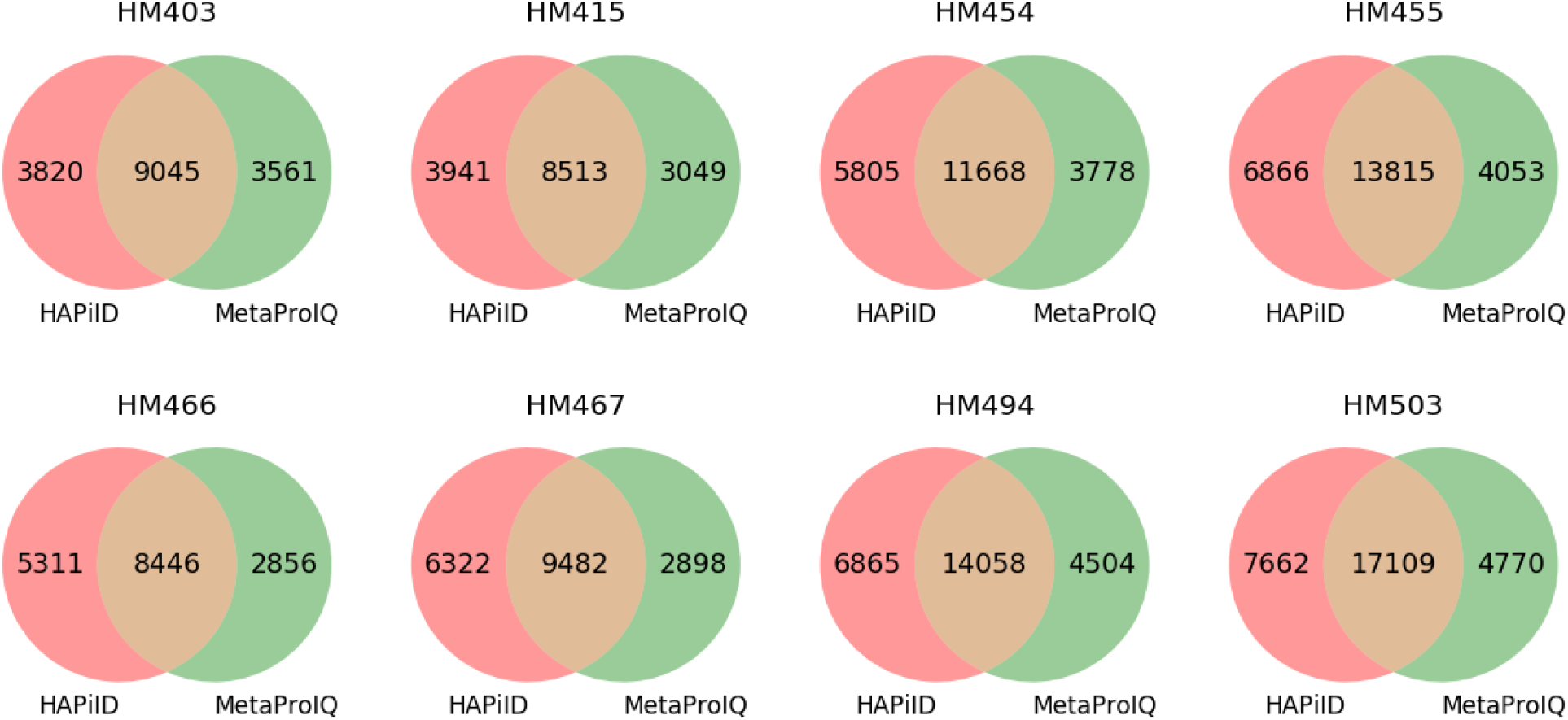
Venn diagram summarizing unique peptides identified by HAPiID and MetaPro-IQ and the overlap between them.

To do so, we downloaded the latest version of the ‘‘Integrated reference catalog of the human gut microbiome” (the IGC-database, which was used by MetaPro-IQ) and then performed a spectra search against this database, to estimate the computation time required to perform the first step in MetaPro-IQ. We note the IGC database contains a total of 9,878,647 genes, more than 8,512,249 protein coding genes predicted from our collection of genomes. Since our proposed method is also composed of two steps and the initial step is used to define the database size for the second (final) step, we compared the database sizes and the running times of each step for the two approaches separately. The results are summarized in Table 3. On average, it took MS-GF+ about 455 CPU hours to complete the spectra search against the IGC database, whereas the first step took HAPiID less than 4 CPU hours to complete. When considering the running time for the whole HAPiID pipeline (both steps), it remains over 50 times faster than the spectra search against the IGC database (which approximated the running time of the first step in MetaPro-IQ pipeline).

**Table 3:**
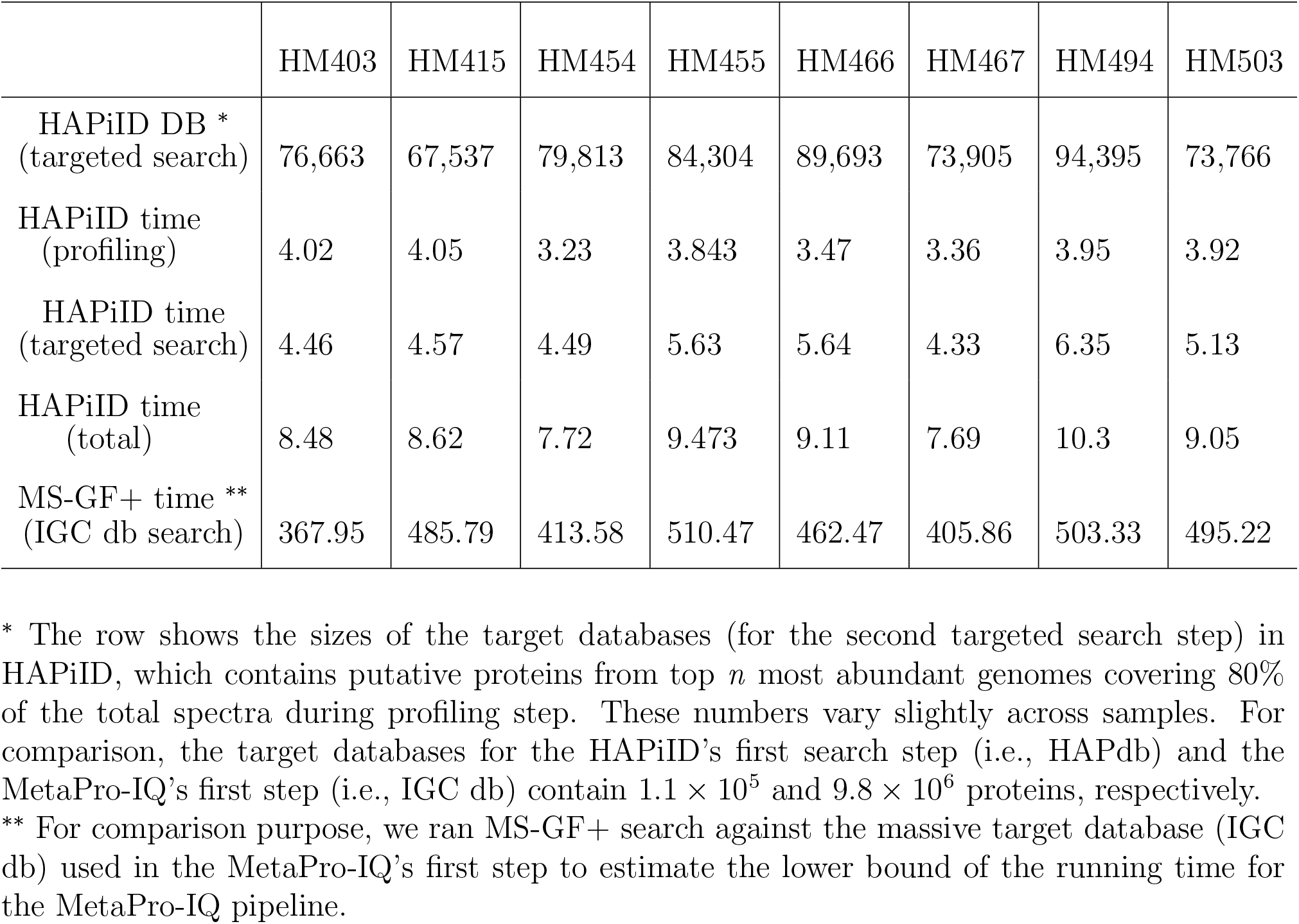
The breakdown of the running time (in CPU hours) for HAPiID.

|  | HM403 | HM415 | HM454 | HM455 | HM466 | HM467 | HM494 | HM503 |
| --- | --- | --- | --- | --- | --- | --- | --- | --- |
| HAPiID DB *<br>(targeted search) | 76,663 | 67,537 | 79,813 | 84,304 | 89,693 | 73,905 | 94,395 | 73,766 |
| HAPiID time<br>(profiling) | 4.02 | 4.05 | 3.23 | 3.843 | 3.47 | 3.36 | 3.95 | 3.92 |
| HAPiID time<br>(targeted search) | 4.46 | 4.57 | 4.49 | 5.63 | 5.64 | 4.33 | 6.35 | 5.13 |
| HAPiID time<br>(total) | 8.48 | 8.62 | 7.72 | 9.473 | 9.11 | 7.69 | 10.3 | 9.05 |
| MS-GF+ time **<br>(IGC db search) | 367.95 | 485.79 | 413.58 | 510.47 | 462.47 | 405.86 | 503.33 | 495.22 |

\* The row shows the sizes of the target databases (for the second targeted search step) in HAPiID, which contains putative proteins from top $n$ most abundant genomes covering 80% of the total spectra during profiling step. These numbers vary slightly across samples. For comparison, the target databases for the HAPiID’s first search step (i.e., HAPdb) and the MetaPro-IQ’s first step (i.e., IGC db) contain $1.1 \times 10^5$ and $9.8 \times 10^6$ proteins, respectively.
\*\* For comparison purpose, we ran MS-GF+ search against the massive target database (IGC db) used in the MetaPro-IQ’s first step to estimate the lower bound of the running time for the MetaPro-IQ pipeline.

### Metaproteomics-based taxonomic profiling of microbial communities

As HAPiID is based on spectral search against proteins predicted from reference genomes or MAGs, once peptides are identified, they can be traced back for estimating the expression of the various species at protein level. Here, we demonstrate this application using case studies of identified peptides from the results of the previous section. Based on the results from HAPiID’s first step (the profiling step), we characterized taxonomic compositions based on the top *N* most prominent species in each sample, that cover 80% of the total number of spectra identified during the first step. Here we quantified taxonomic composition as the number of unique spectra mapped to each of the genomes during the profiling phase. The results are summarized at the *order* level in Figure 5(a). There were a total of 10 orders representing the top *N* species across all eight samples as described above. The top *N* most abundant species represent 2 Phyla, which were *Firmicutes* (45.01%), *Bacteroidota* (54.99 %). It is worth noting that no two samples shared identical species composition at the order level. Individual HM466 contains the most diverse composition with a total of 7 orders, while individual HM503 are the least diverse with a total of 4 orders each.

Furthermore, if we characterize the microbial communities using all the search results from the profiling step, we can get a more comprehensive view of the species present in the different microbial communities. Figure 5(b) shows the clades at the order level present across the different samples with genomes each contributing at least 3 unique peptide hits. Clade diversity at the order level increases by four folds, from 10 clades (based on peptides from all proteins in top *N* genomes) to 40 (based on identified peptides from HAPs of all genomes, limiting to the genomes that have at least three unique peptides identified). The latter clades arise from 14 Phyla, top 5 most abundant ones being *Firmicutes* (53.44%), *Bacteroidota* (42.48 %), (*Proteobacteria*) (1.58%), *Actinobacteriota* (1.52%) and *Cyanobacteria* (1.57%). These compositions were in agreement with previous observations [52, 53]. This diversity increases more than 6 times (to 64 different orders), if we consider all the species having at least one unique peptide being mapped to them, which is summarized in Supplementary Figure S5. These results demonstrate the complexity of the human gut flora reflected even at the proteome level, and reflect on the quantity of the underrepresented species that often appear with very low abundances.

**Figure 5:**
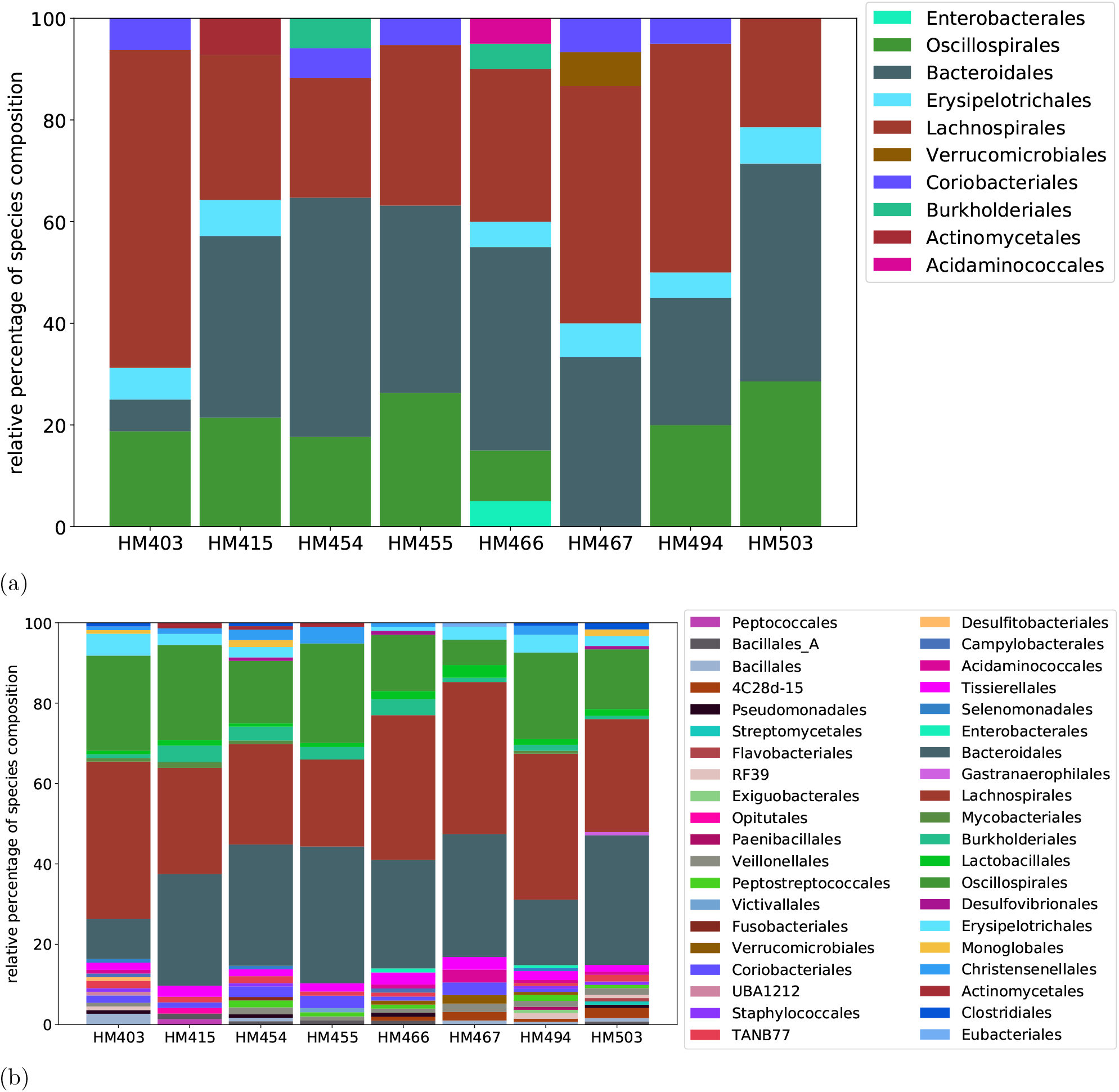
Taxonomic distributions of the eight human gut microbial communities. The distributions are summarized at the order level in the taxonomic hierarchy. (A) shows the distributions considering only the top *N* most abundant species covering 80% of the spectra identified at the profiling state, and (B) shows the distributions using all species each having 3 or more identified spectra based on the results of the profiling step.

### Revealing the functional landscape of abundant species based on metaproteomic data

Although metaproteomics doesn’t provide sufficient data for characterizing proteins from a large number of species in a microbial community, it does provide a fair coverage for the top few most abundant species. So, in addition to providing an overview of what proteins are expressed in microbial communities as a whole, metaproteomics provides opportunities for studying the expression of proteins from individual species, one (or a few) at a time. Table 4 lists the fractions of proteins in the most abundant species that were detected using the metaproteomic data in each sample (and Supplementary Table S4 lists the numbers for the top five most abundant species). Samples HM454, HM455, HM466 and HM467 share the same most abundant species: *Bacteroides vulgatus*, however arising from three different strains. A total of 452 proteins encoded by this species are consistently expressed (containing at least one identified peptide) among samples HM454 and HM466 sharing the same strain. This number decreases to 289 proteins when we only considered those that are supported by at least three spectra. Among the top five genomes that are mostly expressed, on average 19% of their proteins were detected using metaproteomic data with one or more spectra support. This proportion decreases to 10.9% when we restrict proteins supported by at least three or more spectra.

**Table 4:** MS/MS supported proteins in the most expressed species in human gut microbiome.

| Sample | Most abundant species | # putative proteins | #proteins ($\geq 1$ spectrum) all(%) / with-annotation(%) | #proteins ($\geq 3$ spectra) all(%) / with-annotation(%) |
| --- | --- | --- | --- | --- |
| HM403 | *B. xylanisolvens* | 5,442 | 757 (13.9%) / 536 (9.8%) | 437 (8.0%) / 330 (6.0%) |
| HM415 | *B. fragilis* | 5,121 | 751 (14.7%) / 534 (10.4%) | 403 (7.9%) / 313 (6.1%) |
| HM454 | *B. vulgatus* | 4,415 | 740 (16.7%) / 540 (12.2%) | 428 (9.7%) / 346 (7.8%) |
| HM455 | *B. vulgatus* | 4,806 | 1,085 (22.6%) / 744 (15.4%) | 668 (13.9%) / 501 (10.4%) |
| HM466 | *B. vulgatus* | 4,415 | 784 (17.7%) / 583 (13.2%) | 424 (9.6%) / 344 (7.81%) |
| HM467 | *B. vulgatus* | 4,464 | 924 (20.6%) / 659 (14.7%) | 543 (12.1%) / 426 (8.8%) |
| HM494 | *Clostridium_M* | 3,061 | 750 (24.5%) / 556 (18.1%) | 438 (14.3%) / 338 (11.0%) |
| HM503 | *B. ovatus* | 4,931 | 1,077 (21.8%) / 737 (14.9%) | 578 (11.7%) / 438 (8.8%) |

KofamKOALA was able to confidently annotate more than 75% of detected proteins each supported by at least one spectrum, and over 80% of the proteins supported by three or more spectra. The proportion of annotated proteins increased to 93% and 94%, respectively, when we used HMMSCAN and PfamDB to annotate these proteins. This was expected since KofamKOALA uses a much smaller database compared to Pfam to assign proteins to homologous groups.

We grouped the detected proteins in the top five most abundant species into broad functional categories, including Metabolism, Environmental information processing, Organismal systems, Cellular processes, Genetic information processing, Human diseases and uncategorized proteins. Figure 6 shows the relative abundances of the proteins in these functional categories. In general, the protein functional distribution follows similar trends across different samples, with Metabolism being the most abundant category, and Cellular Processes being the least. Supplementary Figures S6 and S7 show the abundance distributions at finer resolution with 48 functional categories. We observed similar trends but were able to see some subtle differences. For example, we saw declined levels of carbohydrate metabolism in HM403 and HM494 compared to the rest. Concerning the Human disease category, the majority of the peptides were mapped to the subcategories including neurodegenerative, endocrine and metabolic and bacterial infectious diseases, with varying relative proportions across samples.

**Figure 6:**
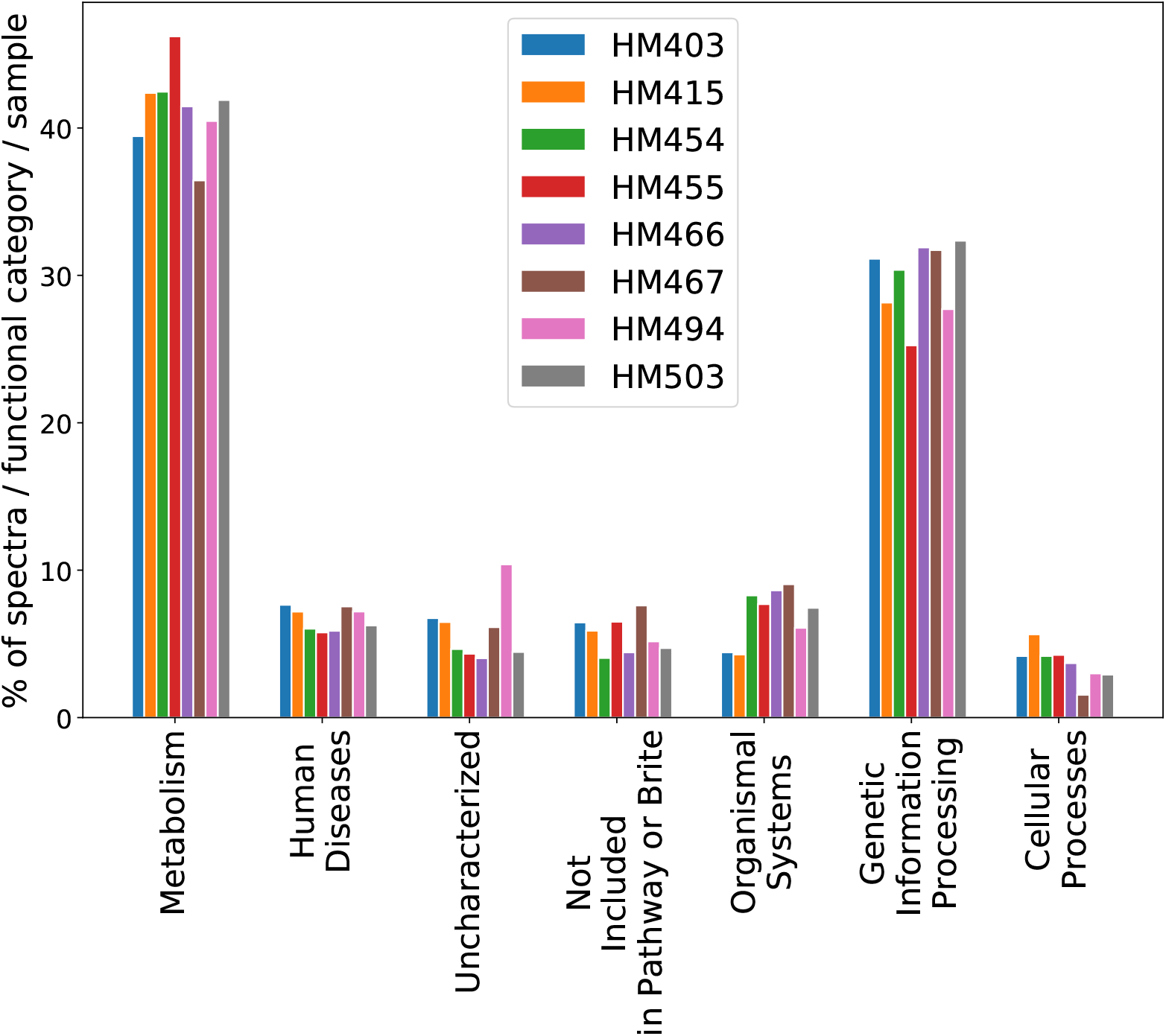
Distribution of identified spectra across the different functional categories.

We note that the most abundant species in all samples belong to *Bacteroides* (but different species or strains), other than individual HM494, whose most abundant species belongs to *Clostridium*. The taxonomic difference between the latter two was reflected at the functional level. The highly expressed functions in *Bacteroides* include *glyceraldehyde 3-phosphate dehydrogenase, phosphoenolpyruvate carboxykinase (ATP), pyruvate-ferredoxin/flavodoxin oxidoreductase, and fructose-bisphosphate aldolase, class II*; on the contrary, the highly expressed functions in *Clostridium* include *formate C-acetyltransferase, glutamate dehydrogenase (NADP+), O-acetylhomoserine (thiol)-lyase and cysteine synthase*, with only *glyceraldehyde 3-phosphate dehydrogenase* common between the two lists.

Finally, we analyzed the genomic context of the genes encoding for the proteins detected in metaproteomics data to check for the presence of structural relationships (i.e. genes located within close proximity or in an operon). As a case study, we selected the most abundant species within sample HM403 and studied the genomic context of its expressed genes. We specifically looked for genes that are found on the same contig, the same strand, that are within 100 bases apart from each other at most and on average have more than 10 spectra supporting their protein products within such a cluster. We identified a total of 25 such clusters satisfying these conditions. All of our identified gene clusters overlapped with the predicted operon structures by *fgenesB*, a Markov chain-based bacterial operon and gene prediction tool [54]. For demonstration purposes, we show the two largest operon structures in this genome that are highly expressed at the protein level, consisting of 23 genes and 5 genes, respectively.

Unsurprisingly, these genes encode for ribosomal proteins including small subunits (*S3, S5, S7, S8, S10, S14, S17 and S19*) and large subunits (*L2, L3, L4, L5, L6, L14, L15, L16, L18, L22, L23, L24 and L30*), and elongation factor (*EF-G*). A total of 387 spectra were matched to these proteins. The second biggest identified operon was another case of functions related to protein translation (large subunits of ribosome, see visualization of these two operons in Figure 7). The other highly expressed operons include genes encoding for transporter proteins, DNA replication machinery, amino acid biosynthesis, starch binding outer membrane protein and pyruvate-ferredoxin/flavodoxin oxidoreductase. See Supplementary Table S5 for all identified operons and their predicted functions.

**Figure 7:**
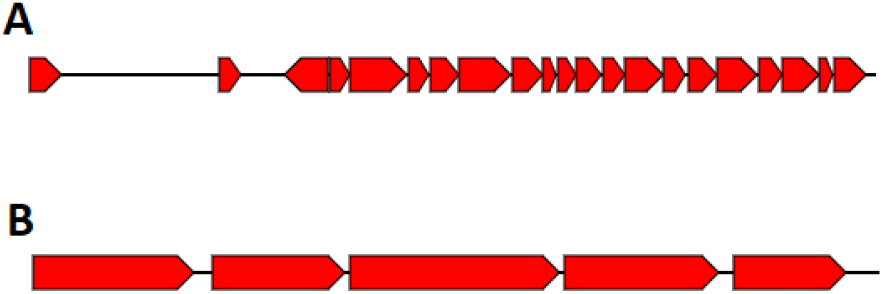
Visualization of the two largest expressed operons in *B. xylanisolvens*. Both operons are found in the contig 25 of its draft genome (*20298_3_31*). Genes are shown as red arrows.

## Discussion

We developed HAPiID, which leverages the HAP guided profiling for creating compact yet effective target database for metaproteomics data analysis. Although the primary goal of developing HAPiID was to speed up the search (by not using the blind search of spectra against a huge database with millions of proteins for peptide identification), the tests showed that HAPiID also achieved significant improvement on peptide identification when compared to MetaPro-IQ [32]. We observed consistent performance improvement of HAPiID using either MG-GF+ or X! Tandem as the search engine. We note that it is possible to further improve the speed of HAPiID by incorporating spectral clustering using our new algorithm msCrush [55], just like MetaLab [33] which adopts PRIDE Cluster [56] for spectra clustering.

HAPiID includes a mechanism to automate the selection of genomes based on the profiling step results to be used in the second step of expanded search: it selects top *n* genomes covering at least 80% of identified spectra from the profiling step. This criterion worked well for the synthetic metaproteomic dataset with low complexity, and also the real gut metaproteomic datasets with higher species complexity. However, this value could be adjusted by the user based on prior knowledge about samples and/or the complexity of the datasets.

In addition to providing a universal target database that can be used across different studies allowing straightforward comparison of the results, HAPiID identifies species that are expressed, rather than providing a list of genes. Thus our pipeline can be utilized to profile a metaproteomic sample by reporting species composition as demonstrated in Results. Such information can be used to further our understanding of the functional contributions of different bacterial species at the proteome level across different samples in various conditions. We annotated functions, as much as possible, using KOFAM and Pfam databases. Characterizing the most abundant protein functions in each sample and each genome allowed us to demonstrate the potential of using a reference based method, such as ours, in revealing functional landscapes across different samples.

It is often a concern that a simple combination of the results from separate spectral searches will underestimate the actual FDR [57]. HAPiID is a two-step approach, however the first step is for profiling, and the final results are only from the second step of expanded search. So the FDR inflation is less of a concern for HAPiID. On the other hand, we introduced the “consistency rate” measure to help us study the impact of using smaller databases for spectral search, and our results show that using smaller databases, as long as they still contain enough genomes, will result in accurate identifications.

We experimented with adding a third step to our pipeline which involves a more focused search over protein sequences that contain considerable number of identified peptides in the second step. By constructing a very small database composed of protein sequences having at least five peptide hits in the second step, we were able to identify on average 10% more unique peptides compared to our two step approach (1,921 additional peptides, see Supplementary Table S6 for more details). However we did not integrate this last step in our final pipeline. Our main concern was the effects of combining the identified peptides from the second step and the new third step over the final FDR value. Each of the steps were controlled to have an FDR of 1% or less; however, combining two steps may result in an actual FDR higher than 1%. Further validations and FDR recalculations would be needed before we can reliably combine results.

For quantification purposes, we used a simple approach based on unique spectra mapped to proteins and in turn genomes to quantify the abundance of the different genomes in samples. While species quantification was not our primary focus in this work, more accurate techniques should be employed that take advantage of the areas under the spectral peaks in order to quantify species from a metaproteomics perspective such as the one used by MaxQuant [58]. In addition, it is worthwhile to consider combining HAPiID with matched-metagenome based approach such as our own Graph2pro/Var2pro [30, 31], to further improve peptide and protein identification from metaproteomic data, when matched metagenome is available.

We note that we focused on human gut metaproteomics in this paper; however, HAPiID can be extended to analyze metaproteomics associated with other hosts (e.g., mouse) or environments, when a good collection of reference genomes/MAGs specific to these microbiomes become available. HAPiID is highly dependent on the initial reference database: peptide identification rate will be greatly affected by the diversity and the quality of the genomes and MAGs included in the database, and incomplete genomes may hinder the ability of our approach to correctly profile metaproteomic samples and select abundant species. With the ongoing progresses of genome/metagenome sequencing, we foresee much broader applications of HAPiID. We include in the HAPiID package scripts for generating reference protein database for spectra match for users who are interested in using HAPiID for different purposes. Finally we note that because HAPiID is a reference-based approach and its efficiency relies on the completeness of the genome collection, a potential pitfall is that it may miss identification of peptides encoded by the accessory genes that are important for understanding the functionality of the underlying microbial communities.

## Conclusions

The HAP based profiling approach provides a novel effective way for guiding the construction of target database for metaproteomic data analysis. Tests of the HAPiID pipeline built upon the HAP profiling approach demonstrated that the pipeline not only drastically reduced the computation time but also improved the peptide identification from spectra data. HAPiID provides a universal approach for analyzing human-gut associated metaproteomic data, facilitating the application of metaproteomics in human microbiome research.

## List of abbreviations

MAG: Metagenome Assembled Genomes
MS: Mass Spectrometry
HAP: Highly Abundant Protein
HAPiID: Highly Abundant Protein guided Metaproteomic IDentification
IBS: Irritable Bowel Syndrome
IBD: Inflammatory Bowel Disease
CDI: Clostridium difficile Infection
FDR: False Discovery Rate
HMP: Human Microbiome Project
HBC: Bacteria Culture Collection
RefSeq: Reference Sequence
UMGS: Unclassified Metagenome Genome Sequences
GTDBTK: Genome Taxonomy DataBase Tool Kit
FGS: Frag Gene Scan
KEGG: Kyoto Encyclopedia of Genes and Genomes
KoFamDB: Kegg Orthology family database
Leu: Leucine
Ile: Isoleucine
IGC: Integrated reference Catalog of the human gut microbiome

## Declarations

### Ethics approval and consent to participate

Not applicable

### Consent for publication

Not applicable

### Availability of data and material

The HAPiID pipeline and all of the required data for running the pipeline are available for download at https://github.com/mgtools/HAPiID.

## Authors’ contribution

All authors contributed equally in designing and performing the experiments, writing and revising the manuscript.

## Acknowledgements

The authors would like to thank Dr. Haixu Tang for helpful discussions about the FDR controls.

