## Supplementary figures and tables for "Using high abundance proteins as guides for fast and effective peptide/protein identification from metaproteomic data"

### Supporting Information

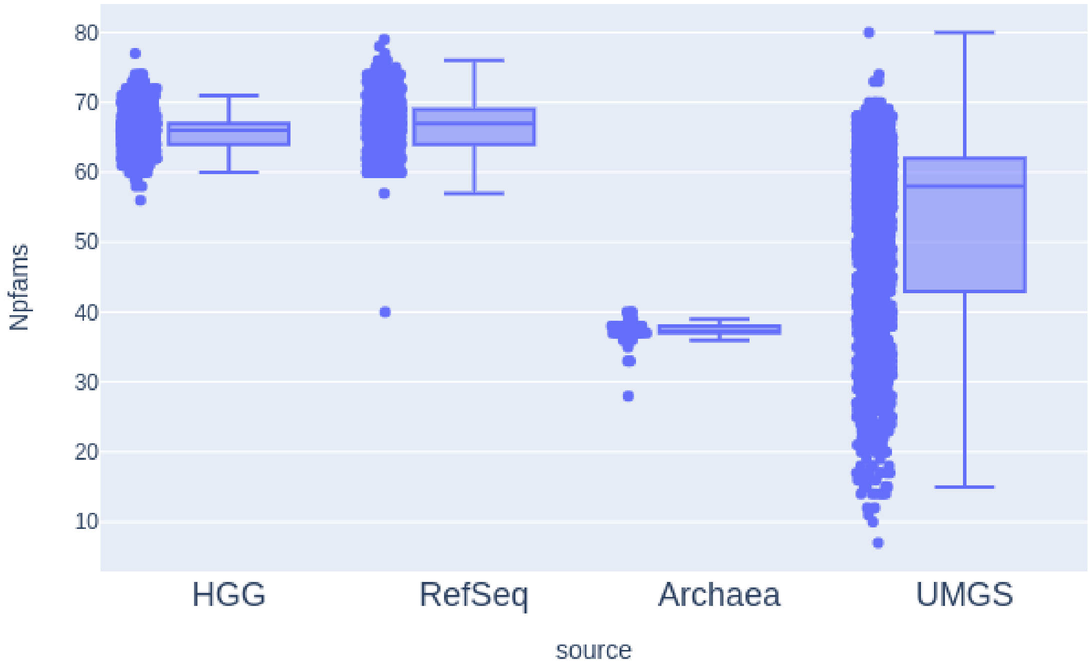


Figure S1: Boxplots summarizing the distribution of the number of marker genes across the different genome sources.

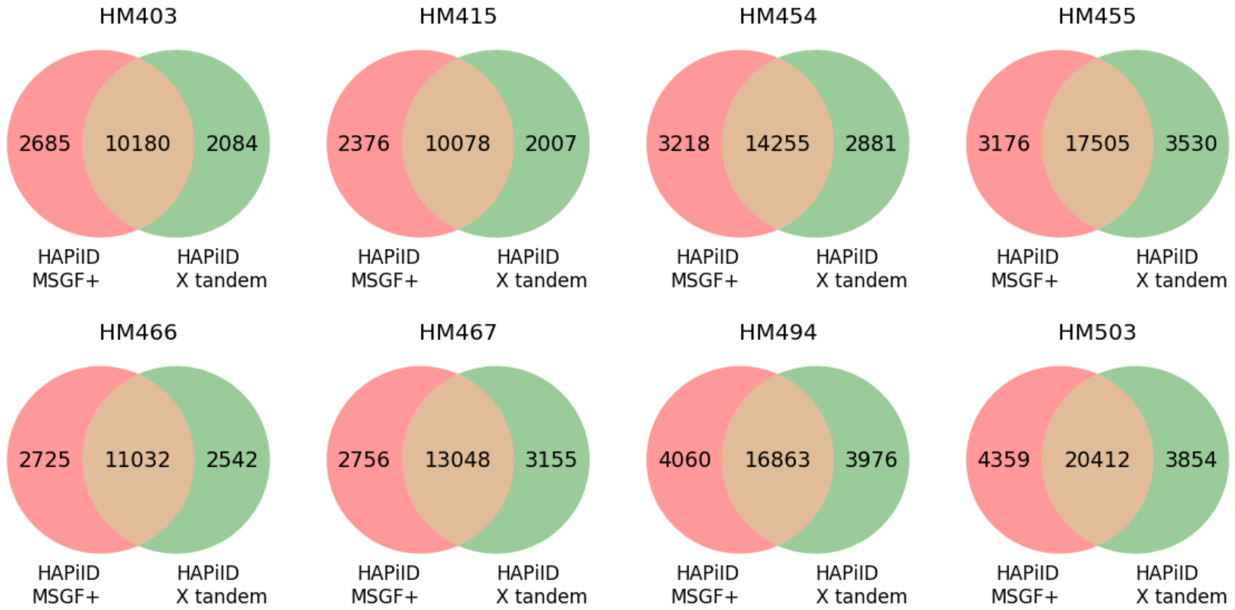


Figure S2: Venn diagram summarizing unique peptide sequences identified by HAPiID using MS-GF+ and HAPiID using X! tandem and the overlap between them.

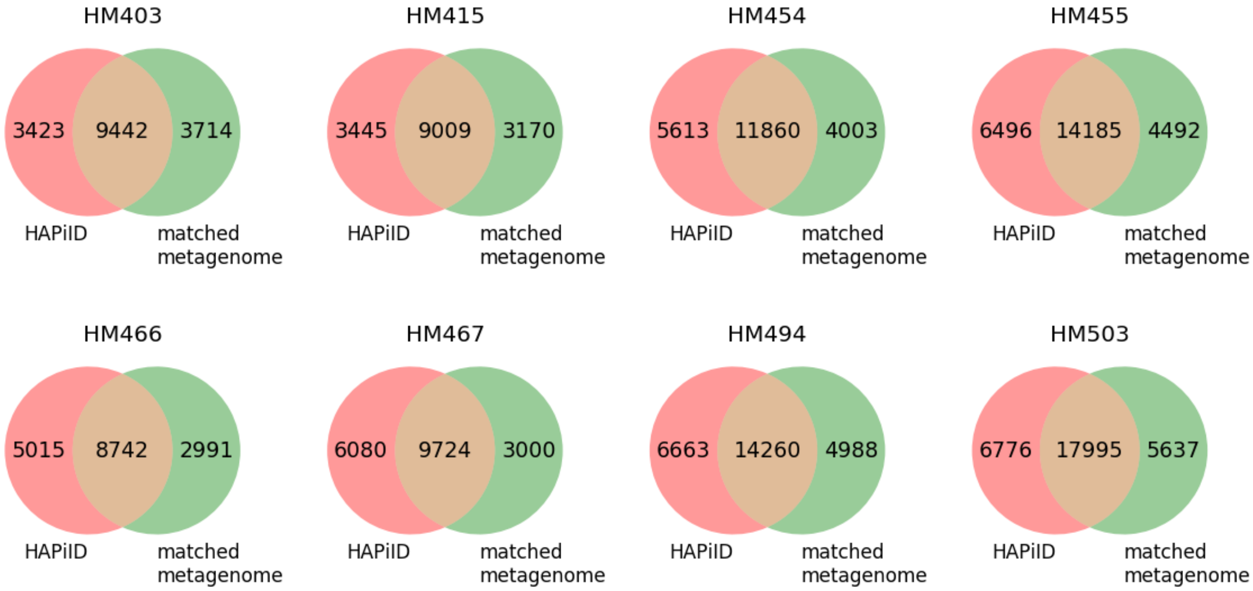


Figure S3: Venn diagram summarizing unique peptides identified by HAPiID and matched metagenome approach and the overlap between them.

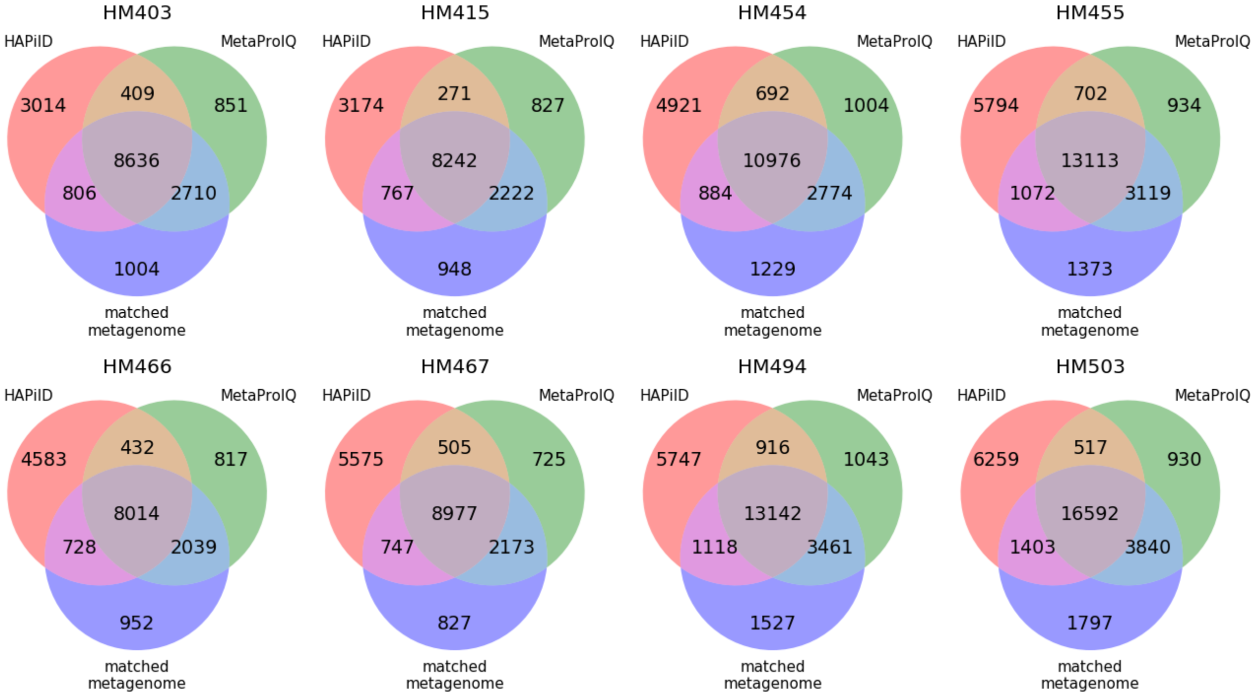


Figure S4: Venn diagram summarizing unique peptides identified by MetaPro-IQ, matched metagenome and HAPiID approach and the overlap between them.

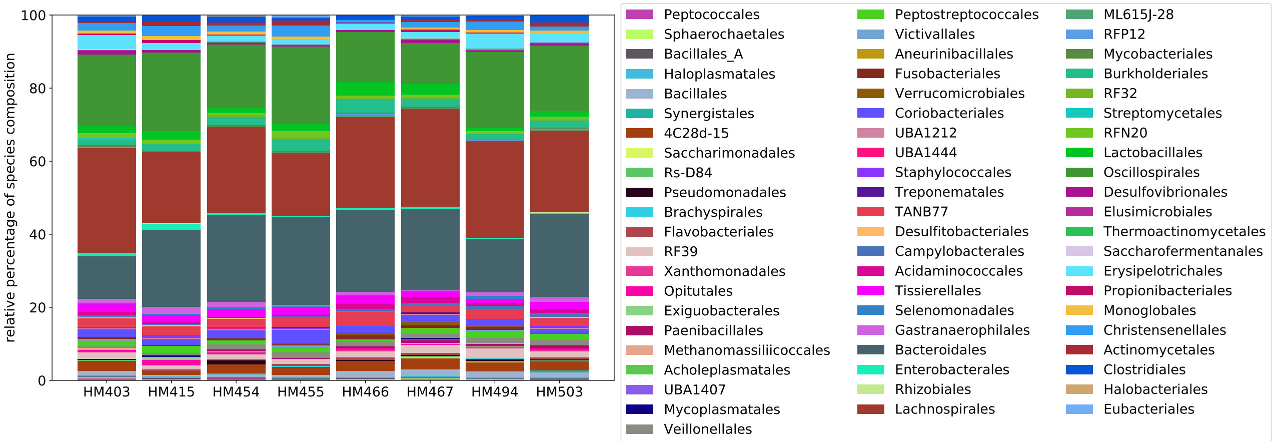


Figure S5: Taxonomic distributions of the eight human gut microbial communities, using all species each having 1 or more identified spectra based on the results of the profiling step.

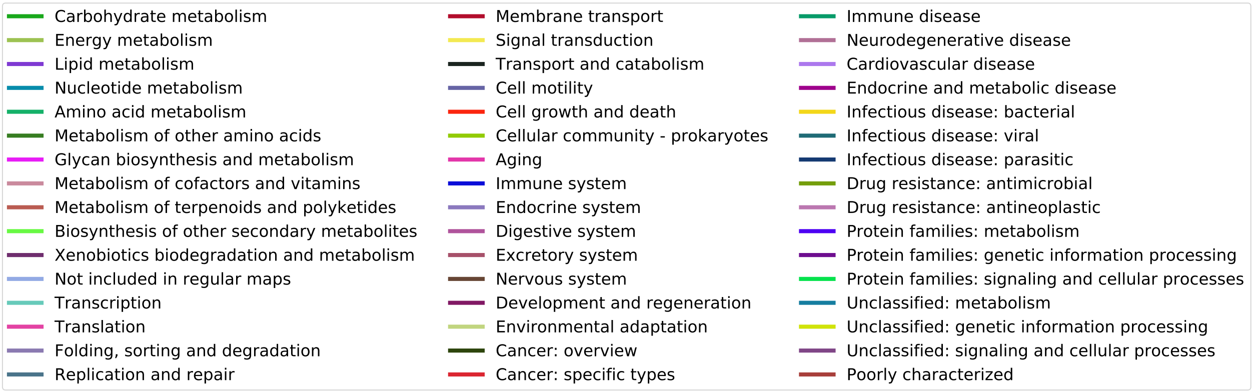


(a)

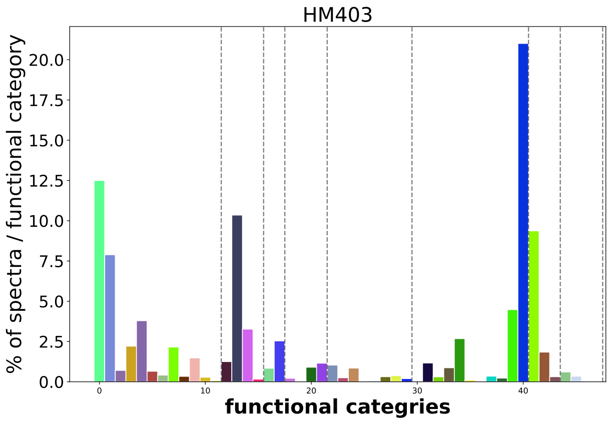


(b)

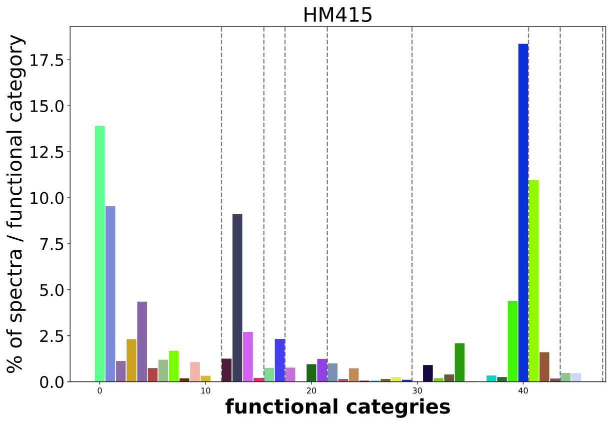


(c)

Figure S6: Distribution of identified spectra across the different functional categories (samples HM403 and HM415).

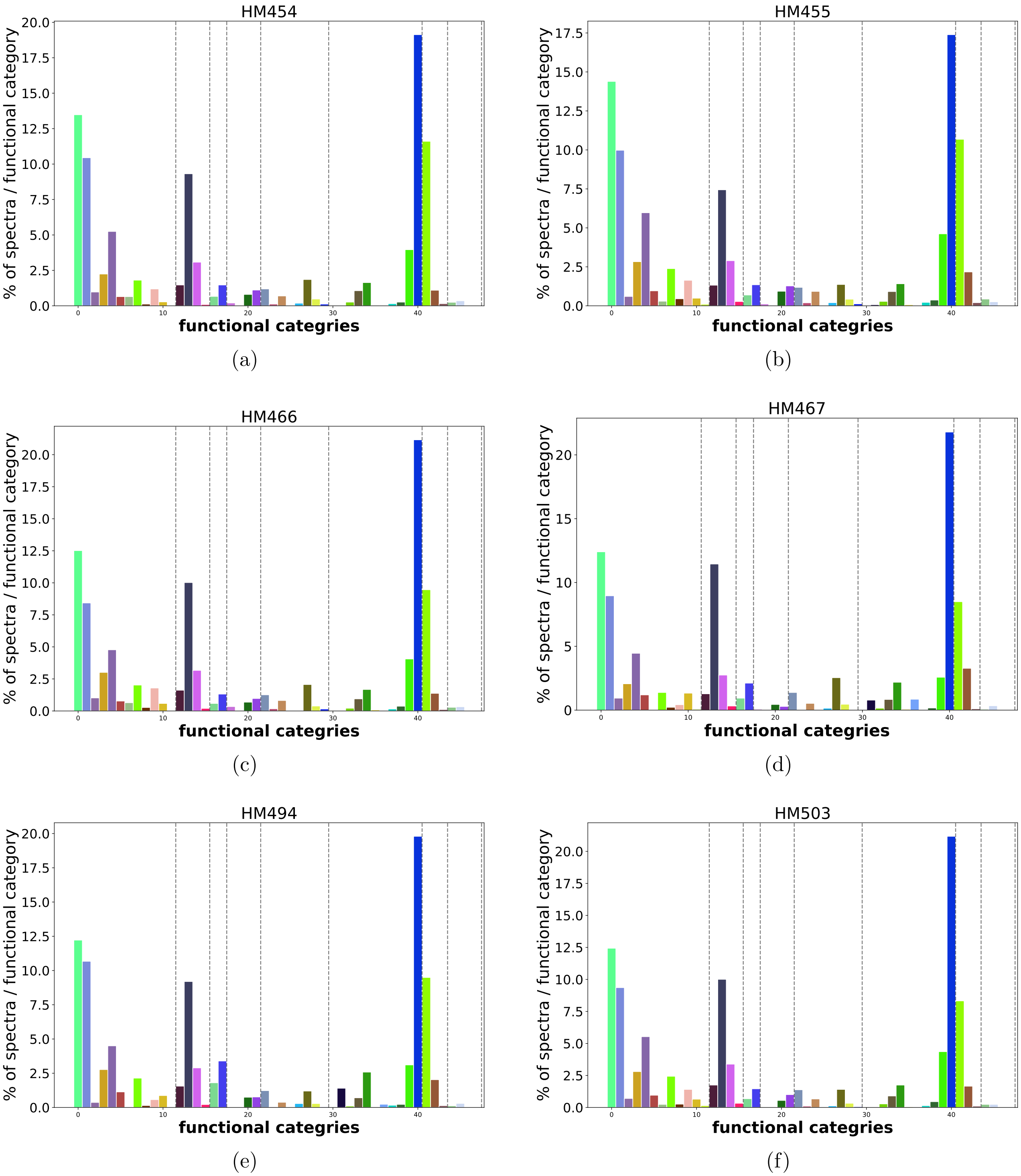


Figure S7: Distribution of identified spectra across the different functional categories (samples HM454, HM455, HM466, HM467, HM494 and HM503).

Table S1: Agreement of identified peptides between searches against target databases of various sizes across samples HM403, HM415, HM454 and HM455

| HM403 DB | top5 | top10 | top20 | top30 | top40 | top50 | top100 |
|---|---|---|---|---|---|---|---|
| top 5 | 100.0 | 99.294 | 98.861 | 98.735 | 98.644 | 98.539 | 98.134 |
| top10 | | 100.0 | 99.475 | 99.343 | 99.25 | 99.156 | 98.787 |
| top20 | | | 100.0 | 99.813 | 99.738 | 99.656 | 99.283 |
| top30 | | | | 100.0 | 99.907 | 99.832 | 99.443 |
| top 40 | | | | | 100.0 | 99.925 | 99.534 |
| top50 | | | | | | 100.0 | 99.629 |
| top100 | | | | | | | 100.0 |
| HM415 DB | top5 | top10 | top20 | top30 | top40 | top50 | top100 |
| top5 | 100.0 | 99.398 | 99.115 | 98.953 | 98.846 | 98.733 | 98.429 |
| top10 | | 100.0 | 99.707 | 99.554 | 99.45 | 99.334 | 99.01 |
| top20 | | | 100.0 | 99.849 | 99.722 | 99.6 | 99.287 |
| top30 | | | | 100.0 | 99.868 | 99.725 | 99.449 |
| top40 | | | | | 100.0 | 99.849 | 99.582 |
| top50 | | | | | | 100.0 | 99.747 |
| top100 | | | | | | | 100.0 |
| HM454 DB | top5 | top10 | top20 | top30 | top40 | top50 | top100 |
| top5 | 100.0 | 99.228 | 98.663 | 98.431 | 98.244 | 98.189 | 97.649 |
| top10 | | 100.0 | 99.453 | 99.231 | 99.03 | 98.968 | 98.461 |
| top20 | | | 100.0 | 99.766 | 99.557 | 99.498 | 99.026 |
| top30 | | | | 100.0 | 99.809 | 99.729 | 99.326 |
| top40 | | | | | 100.0 | 99.921 | 99.509 |
| top50 | | | | | | 100.0 | 99.588 |
| top100 | | | | | | | 100.0 9 |
| HM455 DB | top5 | top10 | top20 | top30 | top40 | top50 | top100 |
| top5 | 100.0 | 99.375 | 98.951 | 98.608 | 98.407 | 98.343 | 97.744 |
| top10 | | 100.0 | 99.523 | 99.158 | 98.961 | 98.883 | 98.344 |
| top20 | | | 100.0 | 99.637 | 99.459 | 99.377 | 98.896 |
| top30 | | | | 100.0 | 99.825 | 99.726 | 99.284 |
| top40 | | | | | 100.0 | 99.914 | 99.479 5 |
| top50 | | | | | | 100.0 | 99.572 |
| top100 | | | | | | | 100.0 |

Table S2: Agreement of identified peptides between searches against target databases of various sizes across samples HM466, HM467, HM494 and HM503

| HM466 DB | top5 | top10 | top20 | top30 | top40 | top50 | top100 |
|---|---|---|---|---|---|---|---|
| top5 | 100.0 | 99.211 | 98.782 | 98.529 | 98.338 | 98.261 | 98.036 |
| top10 | | 100.0 | 99.467 | 99.203 | 99.038 | 98.971 | 98.695 |
| top20 | | | 100.0 | 99.734 | 99.563 | 99.501 | 99.189 |
| top30 | | | | 100.0 | 99.842 | 99.785 | 99.497 |
| top40 | | | | | 100.0 | 99.939 | 99.66 |
| top50 | | | | | | 100.0 | 99.736 |
| top100 | | | | | | | 100.0 |
| HM467 DB | top5 | top10 | top20 | top30 | top40 | top50 | top100 |
| top5 | 100.0 | 99.4 | 99.006 | 98.838 | 98.653 | 98.543 | 98.054 |
| top10 | | 100.0 | 99.545 | 99.395 | 99.245 | 99.134 | 98.718 |
| top20 | | | 100.0 | 99.857 | 99.718 | 99.617 | 99.261 |
| top30 | | | | 100.0 | 99.867 | 99.771 | 99.409 |
| top40 | | | | | 100.0 | 99.903 | 99.597 |
| top50 | | | | | | 100.0 | 99.684 |
| top100 | | | | | | | 100.0 |
| HM494 DB | top5 | top10 | top20 | top30 | top40 | top50 | top100 |
| top5 | 100.0 | 98.933 | 98.008 | 97.688 | 97.48 | 97.296 | 96.673 |
| top10 | | 100.0 | 99.008 | 98.637 | 98.434 | 98.27 | 97.657 |
| top20 | | | 100.0 | 99.601 | 99.375 | 99.191 | 98.629 |
| top30 | | | | 100.0 | 99.76 | 99.558 | 99.026 |
| top40 | | | | | 100.0 | 99.78 | 99.268 |
| top50 | | | | | | 100.0 | 99.48 |
| top100 | | | | | | | 100.0 |
| HM503 DB | top5 | top10 | top20 | top30 | top40 | top50 | top100 |
| top5 | 100.0 | 99.022 | 98.433 | 98.073 | 97.94 | 97.826 | 97.345 |
| top10 | | 100.0 | 99.352 | 98.99 | 98.843 | 98.72 | 98.214 |
| top20 | | | 100.0 | 99.609 | 99.456 | 99.353 | 98.924 |
| top30 | | | | 100.0 | 99.839 | 99.741 | 99.311 |
| top40 | | | | | 100.0 | 99.913 | 99.5267 |
| top50 | | | | | | 100.0 | 99.634 |
| top100 | | | | | | | 100.0 |

Table S3: Comparison of the number of peptides identified by HAPiID (with various database sizes), MetaPro-IQ and matched metagenome approach across the eight human gut metaproteomic datasets

| database | ribP elonF | top5 | top10 | top50 | top100 | top200 | MetaPro-IQ | Matched |
|---|---|---|---|---|---|---|---|---|
| HM403 | 3,170 | 10,602 | 12,014 | 13,099 | 13,328 | 13,409 | 12,617 | 13,165 |
| HM415 | 2,513 | 14,494 | 12,286 | 13,163 | 13,086 | 12,868 | 11,568 | 12,190 |
| HM454 | 3,501 | 10,408 | 16,727 | 18,158 | 17,978 | 17,921 | 15,456 | 15,874 |
| HM455 | 2,881 | 17,081 | 19,156 | 21,079 | 20,933 | 20,684 | 17,879 | 18,695 |
| HM466 | 2,900 | 11,002 | 12,840 | 14,189 | 14,096 | 13,935 | 11,310 | 11,739 |
| HM467 | 2,977 | 14,320 | 15,793 | 16,537 | 16,674 | 16,347 | 12,390 | 12,728 |
| HM494 | 4,049 | 15,246 | 18,752 | 21,339 | 21,659 | 21,115 | 18,571 | 19,255 |
| HM503 | 4,096 | 19,693 | 24,237 | 25,871 | 25,964 | 25,604 | 21,889 | 23,637 |

Table S4: Proteins identified in the top five most expressed species in human gut microbiome.

| Sample | Top 5 species (per sample) | Putative | Supported (≥ 1 spectrum) | Supported (≥ 3 spectra) |
|---|---|---|---|---|
| **HM403** | *Bacteroides xylanisolvens* | 5,442 | 757 ( 536 ) | 437 ( 330 ) |
| | *Faecalibacterium prausnitzii_D* | 2,870 | 363 ( 299 ) | 202 ( 182 ) |
| | *Blautia_A* | 7,973 | 541 ( 460 ) | 236 ( 207 ) |
| | *Faecalibacterium prausnitzii_I* | 1,904 | 370 ( 298 ) | 206 ( 179 ) |
| | *Faecalicatena gnavus* | 3,377 | 360 ( 295 ) | 182 ( 152 ) |
| **HM415** | *Bacteroides fragilis* | 5,121 | 751 ( 534 ) | 403 ( 313 ) |
| | *Bacteroides_B dorei* | 4,678 | 700 ( 529 ) | 396 ( 311 ) |
| | *Bacteroides ovatus* | 5,219 | 634 ( 468 ) | 322 ( 245 ) |
| | *Blautia_A* | 7,973 | 488 ( 399 ) | 215 ( 187 ) |
| | *Faecalicatena gnavus* | 3,434 | 318 ( 251 ) | 151 ( 125 ) |
| **HM454** | *Bacteroides_B vulgatus* | 4,415 | 740 ( 540 ) | 428 ( 346 ) |
| | *Blautia_A* | 7,973 | 842 ( 664 ) | 423 ( 356 ) |
| | *Bacteroides faecis* | 4,819 | 593 ( 442 ) | 310 ( 251 ) |
| | *Parabacteroides merdae* | 3,689 | 563 ( 407 ) | 306 ( 233 ) |
| | *Bacteroides finegoldii* | 3,615 | 496 ( 385 ) | 270 ( 229 ) |
| **HM455** | *Bacteroides_B vulgatus* | 4,806 | 1085 ( 744 ) | 668 ( 501 ) |
| | *Bacteroides stercoris* | 3,305 | 743 ( 553 ) | 431 ( 343 ) |
| | *Parabacteroides distasonis* | 4750 | 719 ( 514 ) | 385 ( 304 ) |
| | *Bacteroides thetaiotaomicron* | 4,959 | 663 ( 510 ) | 316 ( 265 ) |
| | *Blautia_A* | 7973 | 539 ( 442 ) | 207 ( 176 ) |
| **HM466** | *Bacteroide_B vulgatus* | 4,415 | 784 ( 583 ) | 424 ( 344 ) |
| | *Bacteroides fragilis_A* | 48,54 | 590 ( 446 ) | 304 ( 251 ) |
| | *Bacteroides caccae* | 4,418 | 577 ( 448 ) | 285 ( 237 ) |
| | *Bacteroides faecis* | 4,819 | 565 ( 436 ) | 277 ( 227 ) |
| | *Bacteroides_A coprocola* | 3,833 | 493 ( 385 ) | 250 ( 208 ) |
| **HM467** | *Bacteroides_B vulgatus* | 4,464 | 924 ( 659 ) | 543 ( 426 ) |
| | *Bacteroides fragilis* | 4,128 | 691 ( 518 ) | 383 ( 310 ) |
| | *Bacteroides uniformis* | 4,120 | 630 ( 478 ) | 330 ( 269 ) |
| | *Blautia_A* | 7,973 | 547 ( 450 ) | 251 ( 219 ) |
| | *UBA9502* | 2,908 | 428 ( 351 ) | 224 ( 192 ) |
| **HM494** | *Clostridium_M sp000431375* | 3,061 | 750 ( 556 ) | 438 ( 338 ) |
| | *Clostridium_M* | 3,278 | 637 ( 506 ) | 366 ( 308 ) |
| | *Bacteroides caccae* | 4,418 | 732 ( 515 ) | 396 ( 301 ) |
| | *Bacteroides thetaiotaomicron* | 4,959 | 740 ( 525 ) | 383 ( 299 ) |
| | *Blautia_A* | 7,973 | 756 ( 614 ) | 371 ( 326 ) |
| **HM503** | *Bacteroides ovatus* | 4,931 | 1077 ( 737 ) | 578 ( 438 ) |
| | *Bacteroides_A plebeius* | 3,670 | 896 ( 658 ) | 518 ( 416 ) |
| | *Bacteroides_B vulgatus* | 4,415 | 894 ( 657 ) | 503 ( 404 ) |
| | *Faecalicatena gnavus* | 3,434 | 683 ( 533 ) | 390 ( 315 ) |
| | *Bacteroides uniformis* | 3,633 | 727 ( 551 ) | 364 ( 305 ) |

* Numbers in parentheses reflect number of proteins with a confident functional annotation using KofamKOALA.

Table S5: Highly expressed operons in genome (id: 20298_3_31).

| contig | avg Specs | # of genes | start | end | putative functions |
|---|---|---|---|---|---|
| 25 | 16.8 | 23 | 33673 | 45656 | Translation factors<br>Mitochondrial biogenesis<br>Ribosome<br>Ribosome |
| 25 | 28.6 | 5 | 55109 | 57826 | Mitichondrial biogenesis<br>Ribosome biogenesis<br>Ribosome<br>Ribosome |
| 25 | 24 | 3 | 28121 | 30237 | RNA polymerase<br>Transcription machinery<br>DNA repair and<br>recombination proteins<br>Ribosome<br>Ribosome |
| 48 | 30 | 3 | 33634 | 40519 | Transporters |
| 14 | 12 | 3 | 40047 | 41124 | Ribosome<br>Mitochondrial biogenesis |
| 23 | 30.5 | 2 | 83300 | 88113 | Glycolysis<br>Gluconeogenesis<br>Citrate cycle (TCA cycle)<br>Pyruvate metabolism<br>Butanoate metabolism |
| 41 | 16.5 | 2 | 20471 | 23409 | Ribosome biogenesis<br>Purine metabolism<br>Drug metabolism<br>Alanine, aspartate<br>and glutamate metabolism |
| 41 | 11.5 | 2 | 44188 | 47933 | RNA degradation<br>DNA repair and<br>recombination proteins<br>Purine metabolism<br>Drug metabolism - other enzymes<br>and Exosome |
| 73 | 38 | 2 | 5829 | 10436 | Transporters |
| 22 | 10.5 | 2 | 8215 | 12058 | Transporters |
| 38 | 10.5 | 2 | 2496 | 3350 | Ribosome |
| 7 | 21.5 | 2 | 158203 | 160974 | Citrate cycle<br>Butanoate metabolism<br>Oxidative phosphorylation<br>Carbon fixation pathways<br>Citrate cycle<br>Legionellosis |

| | | | | | |
|---|---|---|---|---|---|
| 85 | 21.5 | 2 | 12795 | 14526 | NA |
| 42 | 11 | 2 | 35375 | 37868 | Riboflavin metabolism<br>Folate biosynthesis<br>Quorum sensing<br>Alanine, aspartate and glutamate metabolism<br>Cysteine and methionine metabolism<br>Arginine biosynthesis<br>Arginine and proline metabolism<br>Tyrosine metabolism<br>Phenylalanine metabolism<br>Phenylalanine, tyrosine and tryptophan biosynthesis<br>Isoquinoline alkaloid biosynthesis<br>Tropane, piperidine and pyridine alkaloid biosynthesis<br>Novobiocin biosynthesis<br>Amino acid related enzymes |
| 45 | 16.5 | 2 | 8619 | 12442 | DNA replication proteins<br>DNA repair and recombination proteins |
| 34 | 30 | 2 | 34553 | 36506 | Chaperones and folding catalysts<br>Mitochondrial biogenesis RNA degradation<br>Longevity regulating pathway - worm<br>Type I diabetes mellitus<br>Legionellosis<br>Tuberculosis<br>Messenger RNA biogenesis<br>Chaperones and folding catalysts<br>Mitochondrial biogenesis<br>Exosome |
| 18 | 14.5 | 2 | 31160 | 33681 | NA |
| 18 | 16.0 | 2 | 58680 | 60443 | Pentose phosphate pathway<br>Pentose phosphate pathway<br>Glyoxylate and dicarboxylate metabolism |
| 18 | 13 | 2 | 96037 | 100300 | Starch and sucrose metabolism<br>Necroptosis<br>Biofilm formation - Escherichia coli<br>Insulin signaling pathway<br>Glucagon signaling pathway<br>Insulin resistance |
| 3 | 12 | 2 | 116704 | 119147 | Pentose phosphate pathway<br>Carbon fixation in photosynthetic organisms<br>Biosynthesis of ansamycins<br>Pentose phosphate pathway<br>Fructose and mannose metabolism<br>Carbon fixation in photosynthetic organisms |
| 96 | 13 | 2 | 4632 | 6318 | Enzymes with EC numbers<br>Pentose and glucuronate interconversions |
| 43 | 10.5 | 2 | 19461 | 22129 | Transporters |

| | | | | | |
|---|---|---|---|---|---|
| 15 | 12.5 | 2 | 92081 | 94962 | Pentose and glucuronate interconversions<br>Fructose and mannose metabolism |
| 15 | 12 | 2 | 101383 | 105287 | Biofilm formation<br>Transcription factors<br>Transcription machinery<br>Ribosome biogenesis<br>Aminoacyl-tRNA biosynthesis<br>Amino acid related enzymes<br>Transfer RNA biogenesis |
| 6 | 10 | 2 | 23195 | 26431 | Mitochondrial biogenesis<br>Aminoacyl-tRNA biosynthesis<br>Amino acid related enzymes<br>Transfer RNA biogenesis |

Table S6: Comparison of the number of unique peptides identified by the two steps in HAPiID pipeline and an extra step of searching spectra against a small target database composed of proteins with at least five identified peptide.

| | HM403 | HM415 | HM454 | HM455 | HM466 | HM467 | HM494 | HM503 |
|---|---|---|---|---|---|---|---|---|
| profiling | 3,170 | 2,513 | 3,501 | 2,881 | 2,900 | 2,977 | 4,049 | 4,096 |
| targeted search | 13,149 | 13,220 | 18,087 | 21,009 | 14,257 | 16,602 | 21,492 | 26,011 |
| extra step | 14,301 | 14,607 | 19,930 | 23,411 | 15,723 | 18,537 | 23,876 | 28,810 |
